# Annotation-free sequence-level multimodal graph learning reveals gut microbiome signatures of atherosclerotic cardiovascular disease

**DOI:** 10.64898/2026.05.24.727482

**Authors:** Wenxing Hu, Wenyi Wang, Le Zhang, Yu Vincent Fu

**Author notes:** **To whom correspondence should be addressed:** Le Zhang, Yu Vincent Fu. These authors contributed equally to this work (Wenxing Hu, Wenyi Wang).

## Abstract

**Background:** Atherosclerotic cardiovascular disease (ACVD) remains one of the leading causes of mortality, and the gut microbiome has been implicated in ACVD-related metabolic and inflammatory processes. However, current microbiome-based disease prediction frameworks predominantly rely on annotation-dependent taxonomic or pathway-level abundance profiles. While effective for community-level association analysis, these approaches compress metagenomic information into predefined biological categories and may consequently obscure fine-grained sequence variation, uncharacterized microbial fragments, regulatory sequence signals and sequence-based structure information embedded within metagenomic DNA. Whether disease-relevant microbiome information can be directly learned from raw metagenomic sequences without prior annotation remains incompletely explored.

**Results:** We developed MLMGCN-CVD, an annotation-free sequence-level multimodal graph-learning framework for ACVD prediction directly from gut metagenomic DNA fragments. Rather than relying on taxonomic aggregation, MLMGCN-CVD integrates pretrained genomic language-model embeddings, sequence-derived structural priors, and read-mapping quantitative evidence to learn disease-associated representations at the DNA fragment level. In a primary cohort comprising 218 ACVD patients and 187 healthy controls, MLMGCN-CVD achieved an area under the receiver operating characteristic curve (AUC) of 0.978 under grouped 10-fold cross-validation. External validation on an independent cohort (PRJNA615842) yielded an AUC of 0.940, outperforming relative- abundance-, pathway-, and sequence-only reference frameworks. Importantly, model-prioritized fragments independently recovered multiple microbiome-associated functions previously implicated in ACVD, including trimethylamine (TMA) metabolism, O-antigen of lipopolysaccharide (LPS) biosynthesis, phosphotransferase systems (PTS), and *Enterobacteriaceae*-associated modules. Beyond these established signatures, Rfam-supported analyses identified candidate regulatory RNA-associated signals, including riboswitches and Bacterial small RNAs, suggesting that regulatory sequence information may contribute previously underexplored discriminatory signals within the ACVD gut microbiome. Several representative sequence elements further showed significant correlations with host cardiometabolic indicators.

**Conclusions:** Our findings suggest that microbiome-associated disease signals extend beyond conventional taxonomic or pathway abundance summaries and can be directly inferred from raw metagenomic sequences. By shifting microbiome modelling from annotation-dependent community aggregation toward annotation-free sequence-level representation learning, MLMGCN-CVD provides a framework for uncovering biologically informative regulatory and functional signals embedded within metagenomic data.

## 1 Introduction

Atherosclerotic cardiovascular disease (ACVD) underlies many major cardiovascular events and remains one of the leading contributors to global mortality[1, 2]. Although established clinical risk factors such as dyslipidemia, hypertension, diabetes, and smoking contribute substantially to population-level risk stratification, they only partially explain inter-individual heterogeneity in disease trajectories and provide limited insight into actionable molecular mechanisms[3]. Increasing evidence for the gut-heart axis indicates that gut microorganisms and their metabolites can influence atherogenesis by shaping host inflammation, cholesterol metabolism, and immune homeostasis[4]. Notably, microbial metabolism of dietary choline, phosphatidylcholine, and carnitine can generate pro-atherogenic metabolites such as trimethylamine N-oxide (TMAO), which has been strongly associated with cardiovascular risk in both experimental and clinical studies. Additional microbiome-associated pathways, including phenylacetylglutamine production, bile acid transformation, and endotoxin-related inflammatory signaling, further support the gut microbiome as a mechanistically relevant component of ACVD pathogenesis[5–8].

Driven by advances in shotgun metagenomic sequencing, gut microbiome research has shifted from characterizing taxonomic profiling towards functional characterization and disease prediction[9]. Multiple studies have demonstrated that gut metagenomic features can discriminate ACVD patients from healthy controls, indicating that the gut microbiome contains biologically informative disease-associated signals[10, 11]. Most current microbiome prediction frameworks, however, rely predominantly on engineered features derived from annotation-dependent pipelines, including taxonomic relative abundance, gene abundance, pathway abundance, functional annotations, and microbial co-occurrence networks[12]. While these approaches have improved interpretability and predictive modelling, they fundamentally compress metagenomic information into predefined biological categories determined by reference databases and upstream annotation procedures.

As a consequence, current annotation-dependent microbiome pipelines may systematically discard disease-relevant sequence- level regulatory and structural information prior to downstream modelling. Aggregation into predefined taxonomic or pathway categories can obscure fine-grained sequence variation, uncharacterized microbial fragments, noncoding regulatory elements, and latent structural signals embedded within metagenomic DNA. This information loss is particularly important in disease gut metagenomic studies, where sparse high-dimensional data, cohort heterogeneity, and incomplete reference databases can further weaken robustness and generalizability[13, 14]. Moreover, fragment-level read-mapping evidence, including sequencing depth, coverage, abundance estimates, and copy-number-related features, may provide important quantitative support for sequence-derived signals, yet such information is rarely integrated with sequence-level representations within a unified predictive framework. Collectively, these observations imply that gut microbiome-associated disease information extends beyond conventional abundance summaries and may remain partially embedded within information from raw metagenomic sequences themselves.

Recent advances in genomic foundation models provide a potential solution to this problem. Large-scale self-supervised sequence language models, including DNABERT-derived architectures and the Nucleotide Transformer, have demonstrated the ability to learn transferable contextual and semantic representations directly from raw nucleotide sequences without requiring predefined biological annotations[15–18]. By capturing latent regulatory elements within genomic sequences, these models offer a new opportunity to investigate microbiome-associated disease signals at the sequence level rather than through community-level aggregation alone. At the same time, biologically meaningful sequence signals are inherently multi-scale. Regulatory patterns may depend not only on linear nucleotide composition, but also on local structural tendencies, contextual dependencies, and relationships among neighboring sequence regions. Sequence-derived structural priors, including DNA geometric properties and folding-related conformational features, may therefore provide complementary biological information relevant to regulatory activity[19, 20]. Meanwhile, the secondary structure of RNA made from its primary sequence plays a critical role in different levels of regulation. RNA secondary structures finely regulate RNA fate, including post-transcriptional control, stability, and translational efficiency. Importantly, such single-stranded folding information may provide complementary structural cues for modelling sequence accessibility and local conformational tendencies that could be relevant to ACVD-related regulatory patterns[21, 22]. However, existing metagenomic disease-prediction studies rarely integrate sequence semantics, contextual dependencies, structural priors, and quantitative read-mapping evidence within a unified representation-learning framework.

Because sequence-level microbiome signals are relational and multi-scale, modelling interactions among fragments may be as important as modelling the fragments themselves. Graph-learning strategies provide a natural framework for capturing such relationships by jointly representing local motif-like patterns, long-range contextual dependencies, and topological associations among sequence elements[23, 24]. Nevertheless, graph-based sequence representation learning remains underexplored in microbiome- associated disease prediction, particularly in annotation-free metagenomic settings.

Moreover, translational efforts have been supported by mature microbiome analysis platforms and emerging interpretable learning tools that facilitate statistical analysis, visualization, functional interpretation, and integrative multi-omics exploration[25, 26]. MicrobiomeAnalyst 2.0[27], for example, supports comprehensive statistical, functional, and integrative analysis of microbiome data outputs, whereas recent interpretable frameworks such as MMETHANE further highlight the value of combining predictive modelling with human-readable biological explanations. However, these resources are still largely designed as general-purpose analysis environments and rarely provide end-to-end predictors tailored to specific cardiovascular tasks and grounded in sequence-level modelling[28]. Currently, many metagenomic disease-identification methods are still released primarily as source code or offline workflows, which results in substantial barriers for users without computational expertise, complicating standardized input preparation, result interpretation, and data sharing. Consequently, reproducibility and adoption across teams and application settings remain limited.

To address these challenges, we developed MLMGCN-CVD, an annotation-free multimodal graph-learning framework driven by large-scale sequence language models for ACVD prediction directly from sequence-level gut metagenomic data. Rather than relying on predefined taxonomic or pathway abundance profiles, the framework directly models metagenomic DNA fragments and integrates complementary information from pretrained sequence semantics, long-range contextual representations, and computable structural priors derived from double-stranded geometry and single-stranded folding-related features. These sequence-derived modalities are aligned in a shared latent space and adaptively fused to construct content-aware semantic graphs. In parallel, read-mapping-derived abundance and copy-number features are incorporated as auxiliary global quantitative evidence through a late-fusion branch, allowing the model to combine sequence-level regulatory information with fragment-level read-mapping support. The resulting graphs are processed by a cooperative GCN-CNN backbone to capture both nonlocal dependencies and local discriminative patterns, and complementary double-stranded and single-stranded structure-derived views are further incorporated within a decision-level screening- review scheme rather than simple feature concatenation to improve robustness. Using internal and external metagenomic cohorts, we demonstrate that sequence-level representation learning not only improves predictive performance, but also recovers biologically meaningful microbial functions and identifies candidate regulatory RNA-associated signals linked to cardiometabolic phenotypes. We further implemented MLMGCN-CVD as an accessible web server to support reproducible prediction and downstream application.

This work provides a practical computational framework for microbiome-based ACVD prediction, and suggests that microbiome- associated disease information extends beyond conventional community-level abundance summaries and can be directly inferred from raw metagenomic sequences.

## 2 Methods

### 2.1 Datasets

A high-quality dataset is essential for developing reliable and biologically meaningful predictive models. The primary cohort used in this study was obtained from the landmark study by Jie et al.[11], which included fecal metagenomic shotgun sequencing data from 218 patients with ACVD and 187 healthy controls. The raw sequencing data are publicly available from the European Bioinformatics Institute repository[29] under accession ERP023788. PRJNA615842 was additionally included as an external cohort to evaluate the cross-cohort generalizability of the proposed model.

The preprocessing workflow was designed to generate high-quality microbial sequence fragments and matched read-mapping- derived quantitative features for downstream sequence-level representation learning. For each sample, raw paired-end FASTQ files were first processed using fastp[30] to remove adapter contamination, low-quality reads, reads containing ambiguous bases, and overly short reads. The resulting clean reads were then aligned to the human reference genome hg38 using Bowtie2[31, 32]. Reads not aligned to the human genome were retained as non-human microbial reads for subsequent analysis.

For sequence assembly, non-human microbial reads from each sample were assembled independently using MEGAHIT[33] in a sample-wise de novo assembly manner. The minimum contig length was set to 500 bp, and contigs shorter than 500 bp were excluded from downstream analysis. After assembly, the corresponding non-human microbial reads from each sample were mapped back to its assembled contigs to calculate read-mapping-derived abundance and depth-based copy-number features. These features included mapped-read and mapped-fragment counts, covered bases, coverage breadth, mean sequencing depth, normalized abundance measures including depth per million reads, CPM, RPKM, TPM, and FPKM, and depth-based copy-number proxies, including mean-depth-, median-depth-, and fragment-depth-derived copy-number values together with their log_2_-transformed forms.

The assembled contigs were then standardized into model-compatible sequence fragments. Contigs ranging from 500 to 1000 bp were retained without further splitting, whereas contigs longer than 1000 bp were divided into non-overlapping 1000 bp windows. For fragments derived from a longer parent contig, the read-mapping-derived quantitative features of the corresponding parent contig were assigned to each derived fragment as contig-level sequencing-background information. Missing or infinite values were converted to zero, and the resulting numerical vector was used as a global quantitative feature branch in the late-fusion model. The external cohort was processed using the same preprocessing and feature-generation workflow to ensure consistency with the primary cohort.

To validate cohort grouping and preliminarily assess microbiome differences between ACVD and healthy controls in the primary cohort, we performed principal coordinates analysis and community composition visualization based on a genus-level relative abundance matrix (Fig. 1). PCoA based on Bray-Curtis dissimilarity[34] showed a clear separation trend between ACVD and control samples, with PERMANOVA R² = 0.117 and p = 0.001, indicating significant differences in community structure. This finding is consistent with the original report by Jie et al.[11] and supports the biological distinguishability of the case-control grouping. The genus-level mean relative abundance profiles showed that both groups were dominated by *Bacteroides*, *Prevotella*, *Faecalibacterium*, and *Blautia,* although their relative proportions differed between groups. Overall, these results indicate that ACVD samples differed from controls at the community level, providing a biological basis for subsequent sequence-level multimodal modelling.

**Fig. 1.**
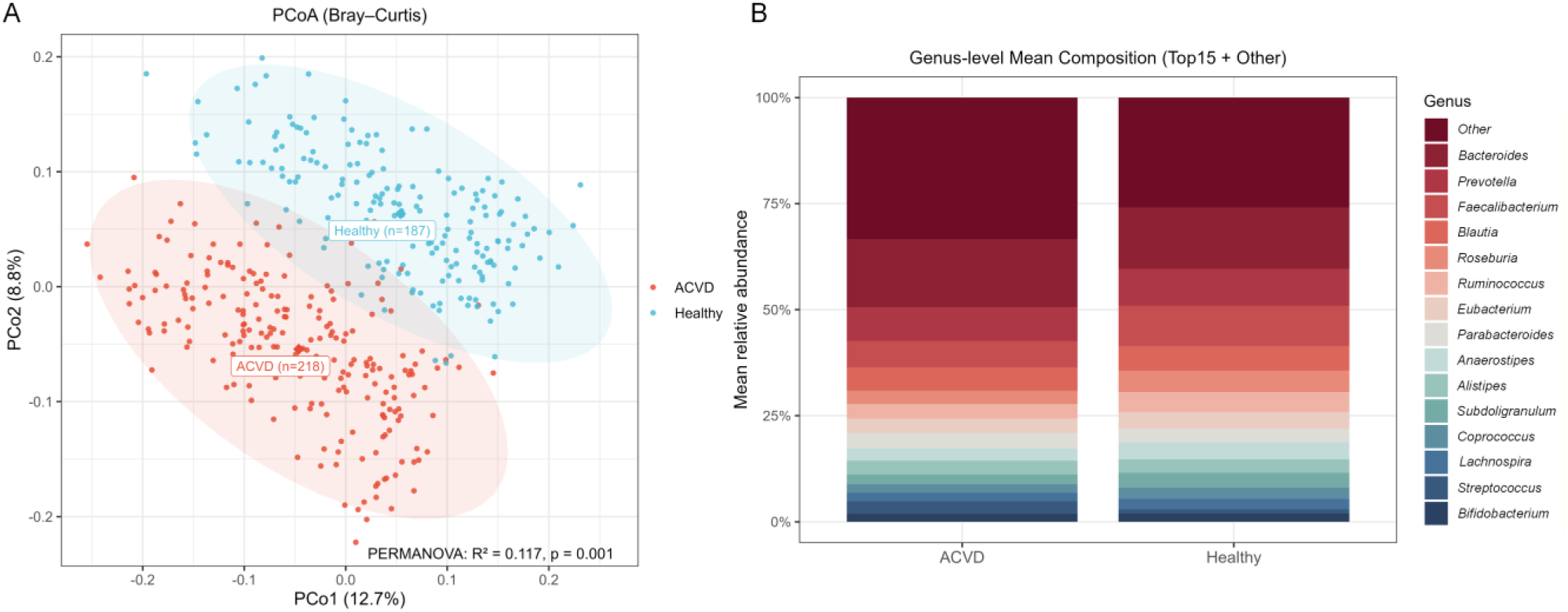
Comparison of gut microbial structure between groups. (A) Principal coordinates analysis based on Bray–Curtis dissimilarity of genus-level relative-abundance profiles from the primary cohort. Each point represents one fecal metagenomic sample, and colors indicate ACVD and healthy control groups. PERMANOVA was used to assess group-level community differences. (B) Mean genus-level relative-abundance composition of ACVD and control samples. The stacked bars show the dominant genera, with low-abundance genera grouped as “Other”.

### 2.2 Framework overview

We propose a multimodal graph learning framework driven by large-scale pretrained sequence language models for gut- microbiome-based ACVD prediction (MLMGCN-CVD), which directly infers ACVD risk from metagenomic DNA fragments (Fig. 2). MLMGCN-CVD derives three complementary token-level sequence representations: (i) sequence semantics from DNABERT- S[35], (ii) long-range context from the Nucleotide Transformer, and (iii) computable structural priors, including dsDNA geometric descriptors from DNAshapeR (e.g., HelT, MGW, ProT, Roll) and ssDNA folding statistics from RNAfold/RNAplfold[36]. These token-level modalities are aligned and adaptively fused into unified node embeddings, which are used to construct a content-aware semantic graph whose edges follow Top-K neighborhoods induced by DNABERT-S attention. In addition, read-mapping-derived abundance and copy-number features are incorporated as auxiliary global quantitative features. Because these features describe fragment-level read-mapping support and copy-number background rather than position-specific token information, they are introduced through a projection-and-gating branch at the late-fusion stage instead of being used for node construction. A Graph Convolutional Network-Convolutional Neural Network (GCN-CNN)[37–43] cooperative backbone then performs multi-scale learning via graph message passing, local convolutional pattern extraction, and a multi-head attention readout to produce fragment-level sequence representations. Finally, the graph-derived fragment representation, the global NT representation, and the projected abundance/copy- number representation are fused before classification. Implementation details are provided in Supplementary Methods S1-S2.

**Fig. 2.**
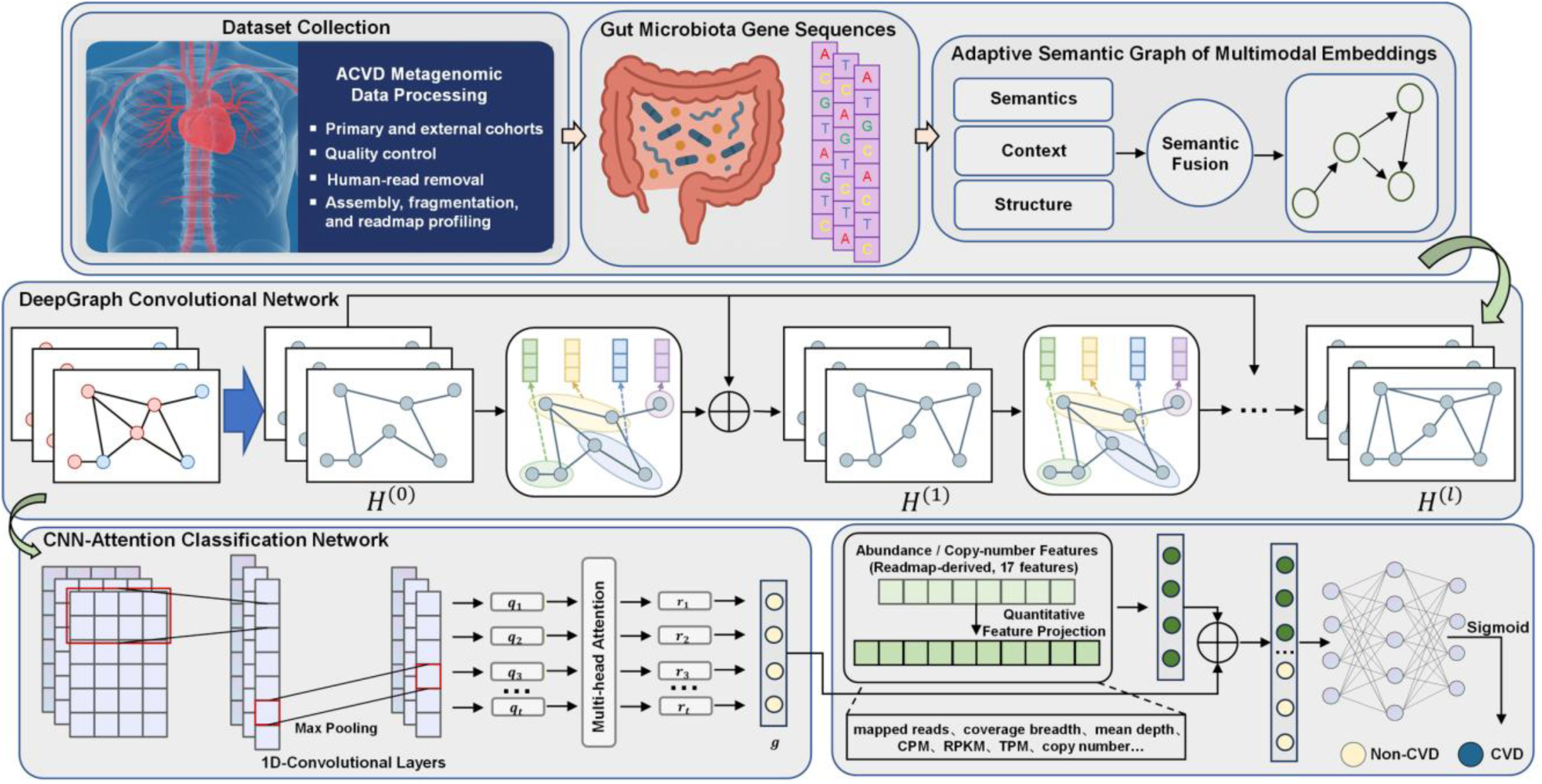
Overall workflow and architecture of MLMGCN-CVD. MLMGCN-CVD processes gut metagenomic reads through quality control, host-read removal, sample-wise assembly, fragment standardization, and read-mapping-derived feature generation. For each standardized DNA fragment, semantic, contextual, and structural representations are adaptively fused to build a content-aware semantic graph. The graph is processed by a cooperative GCN–CNN backbone, and abundance and depth-based copy-number proxies are incorporated through late fusion for fragment-level ACVD classification.

### 2.3 Multimodal representation learning and adaptive fusion

To jointly model semantic information, long-range contextual dependencies, and structural priors, we represent each DNA fragment as a node sequence 𝑉 = [𝑣_1_, …, 𝑣_𝑛_]. Each node corresponds to a 𝑘-mer token of length 𝑘, obtained by a sliding window. For each node 𝑣_𝑖_, we construct three complementary modality-specific representations: semantic, context, and structural modalities. We denote the raw representations of the 𝑖-th node as 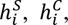 and 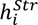. Here, we apply modality-specific projections to align them into a shared latent space detailed by Eq. 1 to address differences in embedding dimensionality and numerical scale across modalities.

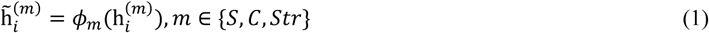

where 𝜙_𝑚_(⋅) is a linear mapping that has a normalization layer to reduce scale differences across modalities and improve representational comparability. Building on this, we employ an adaptive fusion mechanism to dynamically weight the contribution of each modality to the node representations. Specifically, a gating network (Eq. 2) is developed to predict modality weights 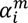 for each node and perform weighted fusion.

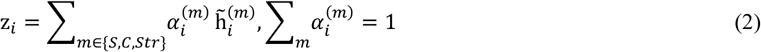

The fused embedding 𝑧_𝑖_ integrates multi-source evidence and serves as the unified input for subsequent semantic-graph construction and graph representation learning. The structural modality is instantiated in two alternative forms: double-stranded geometric priors (dsDNA-Model) and single-stranded secondary-structure priors (ssDNA-Model). For a fair comparison, all components other than the structural channel, including feature extraction, alignment and fusion, and the backbone architecture are kept identical between these two submodules.

### 2.4 Content-aware semantic graph construction

To explicitly model dependencies among nodes within each sequence fragment, we construct a content-aware semantic graph in the fused representation space. For a given sequence fragment with node set 𝑉, the feature vector of node 𝑣_𝑖_ is obtained via multimodal fusion as 𝑧_𝑖_. We determine edges by pairwise node similarity and construct a sparse graph using Top-K nearest neighbors. Specifically, node similarity is defined by Eq. 3.

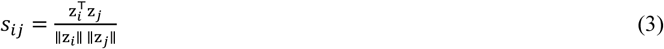

For each node 𝑣_𝑖_, we keep the top 𝐾 most similar neighbors to form 𝑁_𝐾_(𝑖), and we add self-loops to improve propagation stability. The adjacency matrix 𝐴 is defined by Eq. 4.

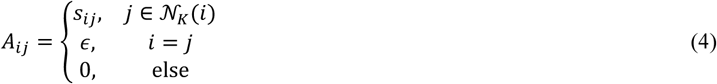

where 𝜖 denotes the self-loop weight, which can be set as a constant or scaled to match the similarity range. To incorporate model-derived priors, we further use DNABERT-S attention to guide neighbor selection and edge weighting, and combine it with the content similarity in Eq. 3, so that the graph reflects both local semantic dependencies and contextual associations. We additionally apply a content-adaptive edge reweighting on top of the fixed K-Nearest-Neighbor (KNN) skeleton to improve robustness under noise and heterogeneity. Finally, each DNA fragment is represented as a content-aware semantic graph 𝐺 = (𝑉, 𝐸, 𝐴), providing an explicit topological prior for subsequent graph learning.

### 2.5 GCN-CNN cooperative backbone and late-fusion classification

Feature-level multimodal fusion alone is insufficient to capture how ACVD-relevant signals are organized, because regulatory cues often consist of long-range dependencies across nodes with local motif and conformational details. We therefore design a GCN-CNN cooperative backbone that jointly models global propagation and local pattern refinement within a unified framework. Given the fused node representation matrix 𝑍 and the content-aware adjacency matrix 𝐴, the GCN branch first performs message passing on the semantic graph to capture long-range dependencies across sequence nodes:

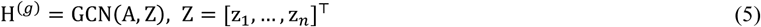

After graph propagation, a 1D convolutional refinement module is applied along the node dimension to enhance local motif-like patterns and short-range structural signals:’

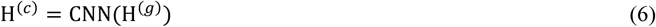

To obtain a graph-derived fragment-level representation, an attention-based readout aggregates the refined node representations into a sequence-level vector:

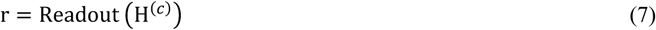

The readout uses learnable query vectors to compute node weights and performs weighted aggregation to emphasize informative regions[44, 45]. In addition to this graph-derived representation, MLMGCN-CVD incorporates two global auxiliary branches before final classification. The sequence-level NT representation is projected into the hidden space to obtain 𝑧_𝑠𝑒𝑞_, and a corresponding gate vector 𝑔𝑎𝑡𝑒 is generated to modulate the graph-derived representation. Meanwhile, the read-mapping-derived abundance/copy-number feature vector 𝑞 is processed by a quantitative feature branch, which produces a projected quantitative representation 𝑧_𝑞𝑢𝑎𝑛𝑡_ and a quantitative gate 𝑞_𝑔𝑎𝑡𝑒_. These abundance and copy-number features describe fragment-level read-mapping support and copy-number background rather than position-specific token information, so they are introduced at the late-fusion stage instead of being used for node construction or semantic graph construction.

The final fused representation is constructed by concatenating the gated graph-derived representation, the projected global NT representation, and the projected quantitative representation:

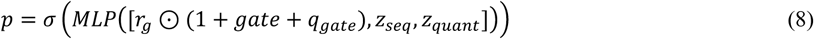

where ⊙ denotes element-wise multiplication. The MLP classifier maps this fused representation to the final ACVD risk probability. We train the model using binary cross-entropy loss[46] and apply regularization and learning-rate scheduling to improve optimization stability. Detailed settings are listed in the Supplementary Information.

### 2.6 Double-stranded and single-stranded submodule

Within the unified MLMGCN-CVD framework, we develop two submodules: one is the dsDNA-Model and the other is ssDNA- Model. Both modules share the semantic modality, contextual modality, semantic graph construction scheme, GCN-CNN backbone, and read-mapping-derived abundance/copy-number branch. They differ only in the structural channel: the dsDNA-Model uses double-stranded geometric features derived from DNAshapeR, whereas the ssDNA-Model uses single-stranded secondary-structure statistics inferred from RNAfold. During inference, we adopt a decision-level two-stage Screening-Review strategy. The dsDNA-Model first outputs a screening probability 𝑝_𝑑𝑠_(𝑥). Fragments with probabilities within the uncertainty interval were considered uncertain and were forwarded to the ssDNA-Model for review. In this study, the uncertainty interval was defined around the decision boundary of 0.5, with 𝜏_𝑙𝑜𝑤_ = 0.5 − 𝛿 and 𝜏_ℎ𝑖𝑔ℎ_ = 0.5 + 𝛿. The default value of 𝛿was set to 0.05, corresponding to an uncertainty interval of 0.45 to 0.55. When fold-specific validation results were available, 𝛿 was selected on the validation split and then applied to the corresponding test fold. Eq. 9 outputs the final probability.

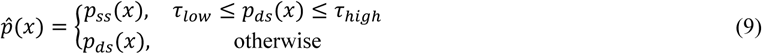

This Screening-Review mechanism avoids noise coupling and dimensionality inflation introduced by feature-level concatenation, while leveraging complementary evidence from double-stranded geometry and single-stranded structure to increase decision stability on borderline fragments and overall predictive robustness.

### 2.7 Experimental settings and evaluation metrics

Model performance was evaluated using sample-grouped stratified 10-fold cross-validation. Although MLMGCN-CVD was trained and evaluated as a fragment-level classifier, cross-validation was grouped by the source individual to prevent information leakage. Specifically, each sequence fragment inherited the ACVD/control label of the individual from which it was assembled. The sample identifier was used as the grouping variable, and the ACVD/control label was used as the stratification variable, following a StratifiedGroupKFold strategy with 10 folds, shuffling enabled, and a random seed of 2020. All fragments derived from the same individual were assigned to the same fold, thereby preventing fragments from the same sample from appearing simultaneously in the training and test sets.

All models were trained for 30 epochs with an initial learning rate of 3 × 10⁻⁴, a weight decay of 1 × 10⁻⁵, and a dropout rate of 0.25. The final backbone used a Post-CNN configuration consisting of six GCN layers followed by two 1D convolutional layers, with a hidden dimension of 768. The batch size was set to 16, and gradient accumulation over two steps was applied to stabilize the effective batch size. We used sensitivity, specificity, accuracy, F1-score, area under the receiver operating characteristic curve, and Matthews correlation coefficient as the evaluation metrics, abbreviated as SN, SP, ACC, F1-score, AUC, and MCC, respectively. These metrics are defined in Eq. 10.

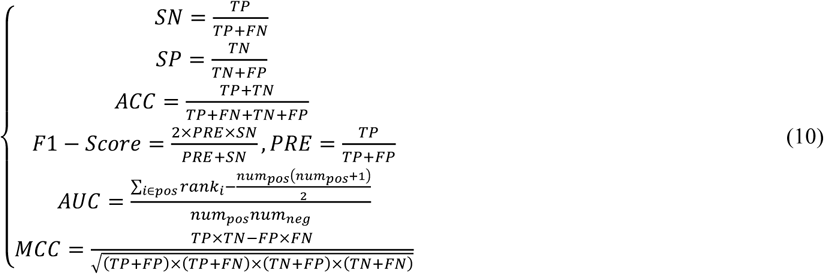

where TP, TN, FP, and FN denote fragment-level true positive, true negative, false positive, and false negative counts, respectively; AUC is the area under the receiver operating characteristic (ROC) curve and measures overall discriminative performance across decision thresholds[47].

## 3 Results

### 3.1 Impact of multimodal sequence embeddings on model performance

To assess whether different sequence-level representations provide complementary evidence for fragment-level ACVD prediction, we systematically compared semantic, contextual, and structural embeddings under single-modality, dual-modality, direct tri-modality concatenation, and adaptive fusion settings. To ensure a controlled comparison, all models were trained and evaluated under the same sample-grouped stratified 10-fold cross-validation protocol, with the model architecture and training configuration kept consistent unless otherwise specified.

Three single-modality baselines were first constructed, namely a DNABERT-S semantic-only model (DNABERT-S-only), a Nucleotide Transformer contextual-only model (NT-only), and a structural-modality-only model (Structural-only). Based on these baselines, we further constructed three dual-modality fusion models, including semantic + structural fusion (Fusion (S+Struct)), structural + contextual fusion (Fusion (Struct+NT)), and semantic + contextual fusion (Fusion (S+NT)), to assess pairwise complementarity among the three representation types. A direct tri-modality concatenation model, denoted as Fusion (All, Concat), was then used as a naive fusion baseline and compared with the proposed adaptive multimodal fusion model (MLMFusion) to determine whether modality alignment and adaptive fusion improved performance beyond simple feature stacking.

As shown in Fig. 3 and Table 1, DNABERT-S-only achieved the highest performance among the single-modality baselines, with mean ACC of 0.816 and F1-score of 0.820. NT-only and Structural-only obtained lower mean ACC values of 0.780 and 0.764, respectively. Pairwise Wilcoxon signed-rank tests showed that DNABERT-S-only significantly outperformed NT-only and Structural- only in ACC and MCC after Benjamini-Hochberg FDR correction, whereas the difference between NT-only and Structural-only was not significant.

**Fig. 3.**
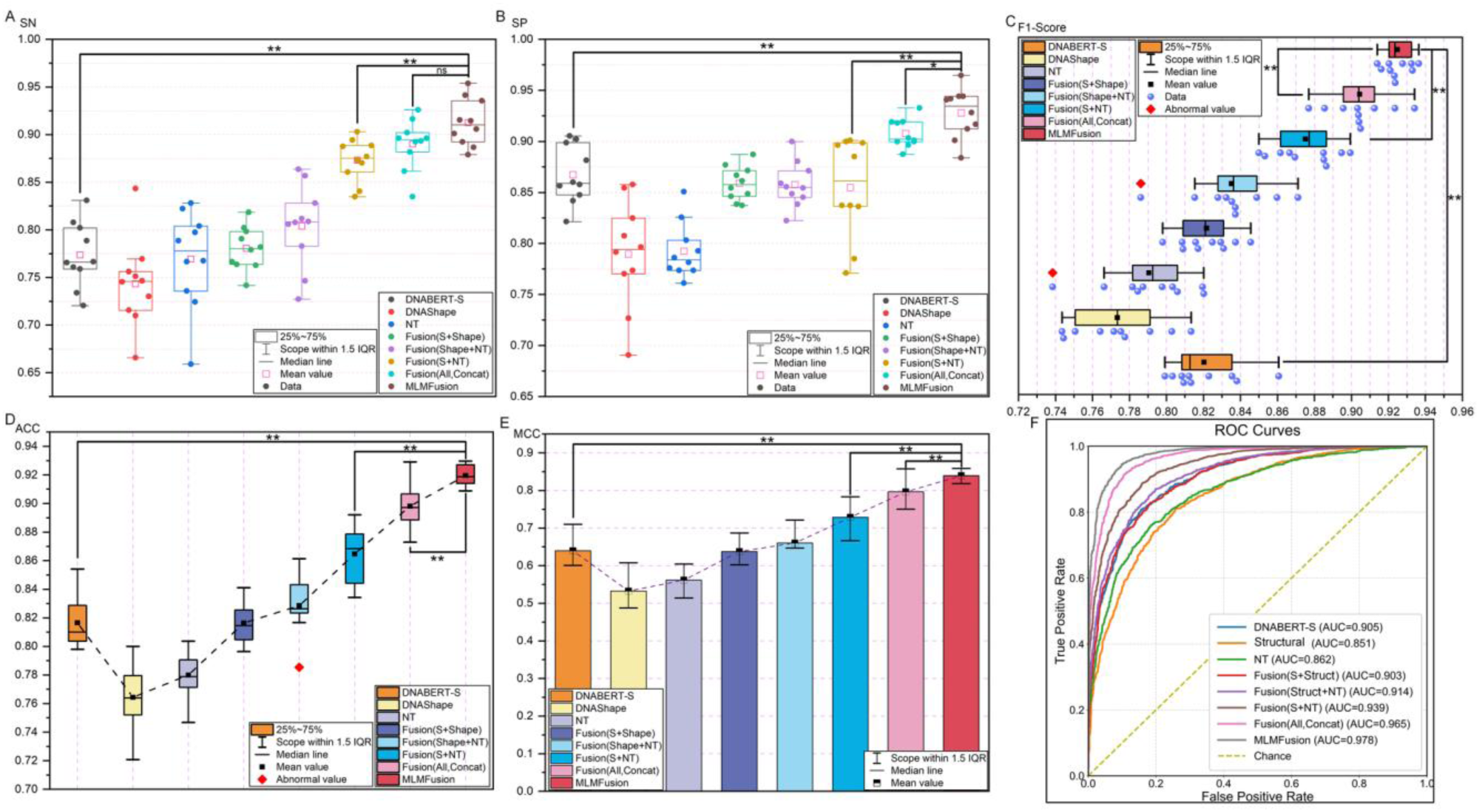
Comparison of model performance across different feature-mode combinations. (A–E) Fold-wise distributions of SN, SP, F1-score, ACC, and MCC for single-modality baselines, dual-modality fusion models, tri-modality direct concatenation, and the proposed adaptive multimodal fusion model. The compared feature modes include DNABERT-S semantic embeddings, Nucleotide Transformer contextual embeddings, and sequence-derived structural features. (F) ROC curves of the compared models. All models were trained and evaluated using the same sample-grouped stratified 10-fold cross-validation protocol. S denotes DNABERT-S semantic features, NT denotes Nucleotide Transformer contextual features, and Structural denotes sequence-derived structural features. Significance annotations, where shown, are based on paired Wilcoxon signed-rank tests with Benjamini–Hochberg correction across predefined comparisons.

**Table 1.** Model Performance Averages from 10-fold Cross-Validation Across Different Feature Modes.

| Metric | DNABERT-S | NT | Structural | Fusion (S+Struct) | Fusion (Struct+NT) | Fusion (S+NT) | Fusion (All, Concat) | MLMFusion |
| --- | --- | --- | --- | --- | --- | --- | --- | --- |
| ACC | 0.816 | 0.780 | 0.764 | 0.816 | 0.828 | 0.865 | 0.898 | 0.920 |
| MCC | 0.640 | 0.562 | 0.533 | 0.638 | 0.660 | 0.729 | 0.796 | 0.840 |
| AUC | 0.905 | 0.862 | 0.851 | 0.903 | 0.914 | 0.939 | 0.965 | 0.978 |
| F1-Score | 0.820 | 0.790 | 0.773 | 0.822 | 0.835 | 0.875 | 0.904 | 0.925 |
| SN | 0.774 | 0.769 | 0.743 | 0.781 | 0.804 | 0.873 | 0.89 | 0.913 |
| SP | 0.867 | 0.792 | 0.789 | 0.859 | 0.858 | 0.855 | 0.908 | 0.928 |

Among the dual-modality models, Fusion (S+NT) achieved the highest overall performance, with mean ACC of 0.865, MCC of 0.729, AUC of 0.939 and F1-score of 0.875. Fusion (S+Struct) and Fusion (Struct+NT) obtained mean ACC values of 0.816 and 0.828, respectively. The tri-modality concatenation baseline further increased performance, reaching mean ACC of 0.898, AUC of 0.965 and F1-score of 0.904.

MLMFusion achieved the best performance across all compared feature-mode settings, with mean ACC of 0.920, MCC of 0.840, F1-score of 0.925 and AUC of 0.978. The ROC curves in Fig. 3F showed the highest AUC for MLMFusion. Supplementary Table S1 showed that MLMFusion significantly outperformed Fusion (S+Struct), Fusion (Struct+NT), Fusion (S+NT) and Fusion (All, Concat) in ACC and MCC after Benjamini-Hochberg FDR correction. Compared with Fusion (All, Concat), MLMFusion showed Δmean values of 0.0214 for ACC and 0.0433 for MCC. In addition, supplementary comparisons against RF, LR, KNN, SVM, and XGBoost under the same MLMFusion representation are reported in Supplementary Results S1.

### 3.2 Impact of cooperative backbone on model performance

To evaluate how cooperative backbone design and dual-view structural inference affect fragment-level ACVD prediction, we performed two sets of controlled experiments under the same training configuration. The first experiment assessed whether the ordering of graph propagation and convolutional refinement influenced model performance by comparing Pre-CNN and Post-CNN backbones across different CNN-layer and GNN-layer settings. This analysis was designed to determine whether nonlocal graph-based dependency modelling should be performed before or after local convolutional pattern refinement. The second experiment compared the standalone dsDNA-Model, the standalone ssDNA-Model, and the proposed Screening-Review strategy to assess whether double- stranded geometric features and RNAfold-derived single-stranded folding-related priors provided complementary evidence during inference.

#### 3.2.1 Ablation on convolution placement and network depth

To examine how local convolutional refinement should be coordinated with graph propagation, we compared two cooperative backbone variants. In the Pre-CNN setting, the convolution module was applied before graph convolution, whereas in the Post-CNN setting, graph propagation was performed before convolutional refinement. For both variants, the numbers of CNN layers and GNN layers were varied to generate different depth combinations. The input features, graph construction scheme, and training hyperparameters were kept unchanged across all compared settings.

Fig. 4 shows the performance of the two backbone variants across different depth combinations. Post-CNN achieved higher performance than Pre-CNN across most CNN-layer and GNN-layer settings. Models with moderate CNN and GNN depths generally performed better than zero-layer or excessively deep configurations. Among all tested settings, the Post-CNN model with two CNN layers and six GNN layers achieved the highest overall performance across ACC, AUC, F1-score and MCC.

**Fig. 4.**
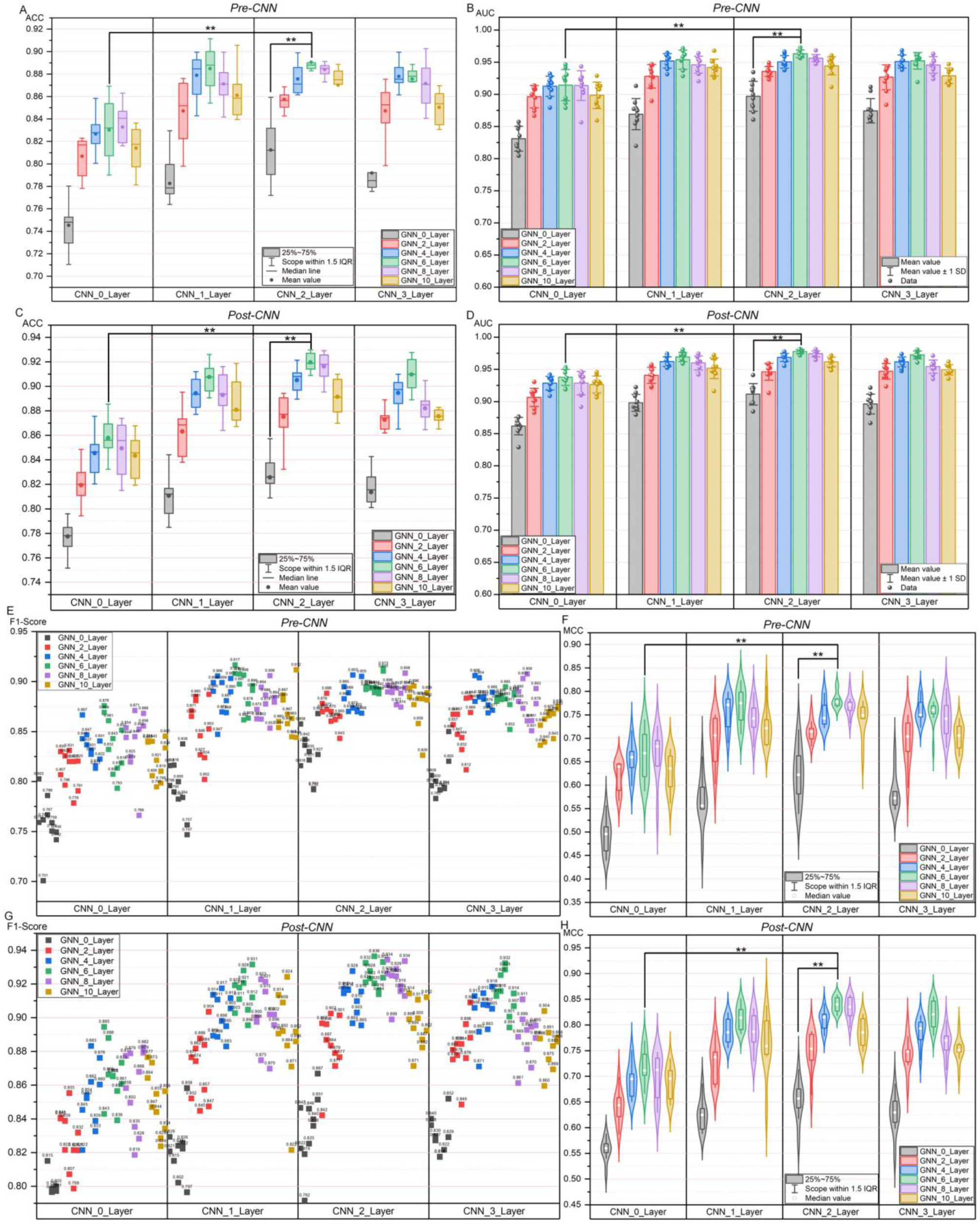
Ablation results of convolution placement and network depth. (A) ACC and (B) AUC of Pre-CNN; (C) ACC and (D) AUC of Post-CNN; (E) F1-score and (F) MCC of Pre-CNN; (G) F1-score and (H) MCC of Post-CNN under different CNN-layer and GNN-layer settings.

The paired Wilcoxon signed-rank tests in Supplementary Table S2 showed that Post-CNN significantly outperformed Pre-CNN in ACC and MCC after Benjamini-Hochberg FDR correction across averaged depth comparisons. Based on these results, the Post- CNN configuration with CNN_2_Layer and GNN_6_Layer was used as the final backbone setting.

#### 3.2.2 Comparison of dsDNA/ssDNA sub-models and Screening-Review strategy

We further evaluated whether the two structural views contributed complementary predictive information. The dsDNA-Model and ssDNA-Model shared the same semantic modality, contextual modality, semantic graph construction strategy, GCN-CNN backbone, and read-mapping-derived quantitative branch. They differed only in the structural channel: the dsDNA-Model used double- stranded DNA shape features, whereas the ssDNA-Model used RNAfold-derived single-stranded folding-related features. Under the same training configuration and data-splitting protocol, three inference settings were compared: the dsDNA-Model alone, the ssDNA- Model alone, and the proposed Screening-Review strategy.

As shown in Fig. 5A, all three settings achieved stable performance across ACC, AUC, MCC and F1-score. The ssDNA-Model obtained slightly higher AUC than the dsDNA-Model. The Screening-Review strategy achieved the highest overall performance among the three settings. The ROC curves in Fig. 5B showed AUC values of 0.965 for dsDNA-Model, 0.967 for ssDNA-Model and 0.978 for Screening-Review.

**Fig. 5.**
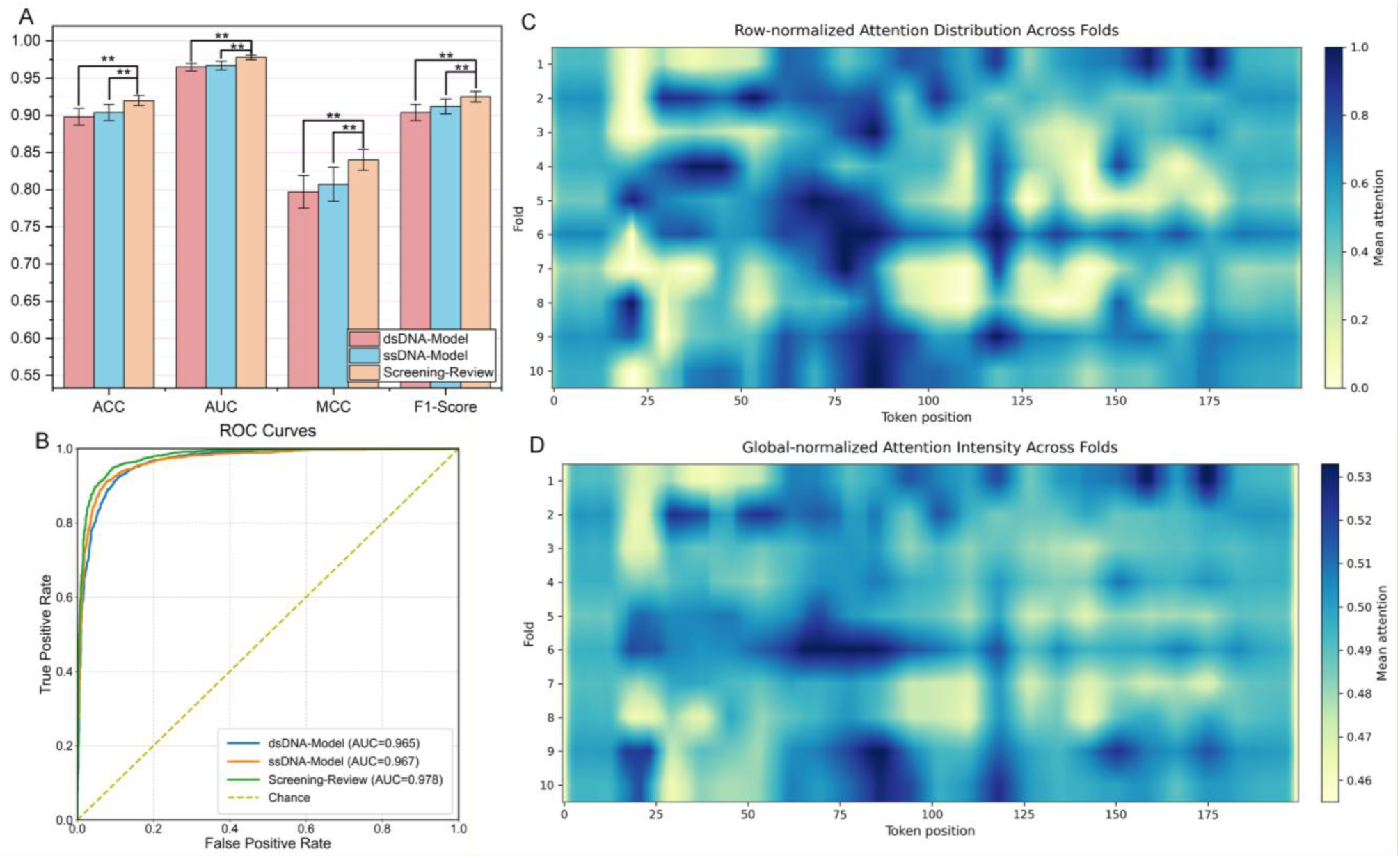
Comparison of dual-view structural models and the Screening-Review strategy. (A) Performance comparison of the dsDNA-Model, ssDNA-Model, and the proposed Screening-Review strategy across ACC, AUC, MCC, and F1-score. The dsDNA-Model uses double-stranded DNA shape features, whereas the ssDNA-Model uses RNAfold-derived single-stranded folding-related features. (B) ROC curves of the three inference settings. (C) Row-normalized attention distribution across cross-validation folds, showing relative attention allocation within each fold. (D) Global-normalized attention intensity across folds, showing the overall distribution of high-attention regions.

Supplementary Table S3 showed that the difference between the dsDNA-Model and ssDNA-Model was not significant in ACC or AUC after Benjamini-Hochberg FDR correction. In contrast, Screening-Review significantly outperformed the dsDNA-Model in ACC and AUC, with Δmean values of 0.0212 and 0.0130, respectively. It also significantly outperformed the ssDNA-Model, with Δmean values of 0.0158 for ACC and 0.0102 for AUC.

These results suggest that neither the dsDNA nor the ssDNA structural view was uniformly dominant, whereas their decision- level collaboration improved predictive performance. Rather than directly concatenating the two structural feature views, Screening- Review introduced the ssDNA branch only for uncertain dsDNA predictions. This design reduced unnecessary noise coupling while allowing additional folding-related evidence to refine borderline predictions. The attention maps in Fig. 5C and Fig. 5D showed nonuniform attention allocation across folds, indicating that the model concentrated decision weights on a subset of informative sequence regions.

### 3.3 External-cohort evaluation on PRJNA615842

To further evaluate the cross-cohort generalizability of MLMGCN-CVD, we performed external-cohort evaluation on the PRJNA615842 dataset. The external cohort was processed using the same preprocessing and feature-generation workflow as the primary cohort, including quality control, host-read removal, sample-wise assembly, fragment standardization, and read-mapping- derived quantitative feature generation. The trained models were applied to PRJNA615842 without retraining or parameter updating. To benchmark external-cohort performance, MLMGCN-CVD was compared with three representative reference methods: Baseline- Relative-Abundance, Baseline-Pathway, and Sequence-Only. These reference methods represent relative-abundance-based community composition, pathway-level functional representation, and sequence-only modelling without multimodal structural integration, respectively. Consistent with the main evaluation setting, external-cohort metrics were calculated at the fragment level unless otherwise specified. In addition to standard classification metrics, stage-wise UMAP visualization was performed to qualitatively examine how the learned feature space was reorganized during graph learning and convolutional refinement.

As shown in Fig. 6A-C, MLMGCN-CVD achieved the highest overall performance among the compared methods in ACC, MCC and F1-score. The ROC curves in Fig. 6D showed that MLMGCN-CVD achieved an AUC of 0.940, compared with 0.925 for Baseline- Pathway, 0.904 for Sequence-Only and 0.903 for Baseline-Relative-Abundance. Detailed statistical comparisons are reported in Supplementary Table S4. These results indicate that MLMGCN-CVD retained discriminative performance in an independent cohort and outperformed abundance-based, pathway-based, and sequence-only reference frameworks under the external-cohort setting.

**Fig. 6.**
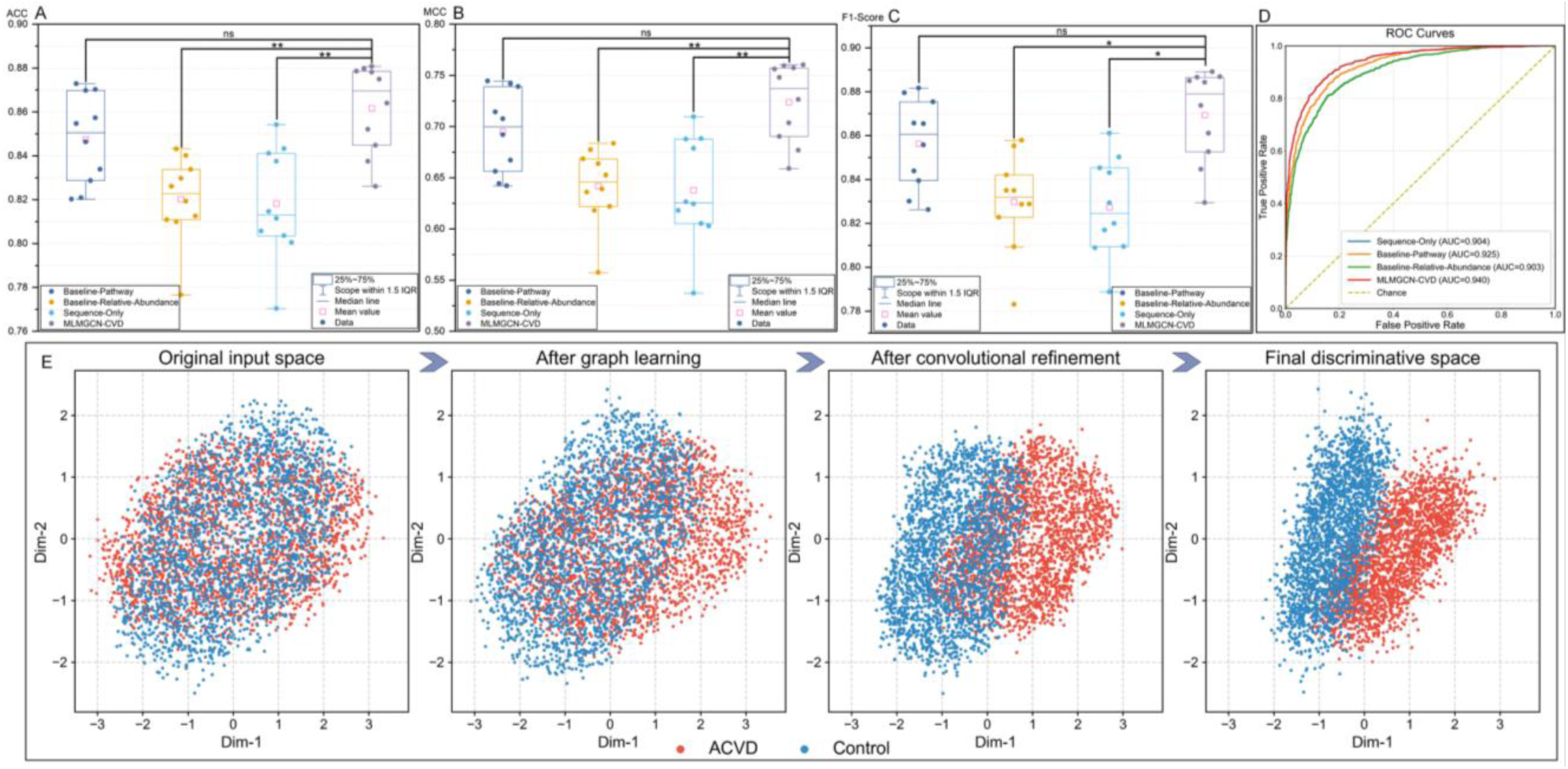
External-cohort evaluation results on PRJNA615842. (A–C) ACC, MCC, and F1-score distributions of MLMGCN-CVD and reference methods on the external cohort. The reference methods include relative-abundance-based, pathway-based, and sequence-only baselines. (D) ROC curves of the compared methods on PRJNA615842. (E) Stage-wise UMAP visualization of the learned feature space in MLMGCN-CVD, showing the original input space, the representation after graph learning, the representation after convolutional refinement, and the final discriminative space.

The stage-wise UMAP visualization in Fig. 6E showed progressive representation changes of MLMGCN-CVD on PRJNA615842. ACVD and control fragments showed substantial overlap in the original input space. After graph learning, the two classes showed clearer organization, and class separation increased further after convolutional refinement. The final discriminative space showed the clearest separation trend. This visualization suggests that graph learning and convolutional refinement progressively reorganized the feature space, but the UMAP analysis should be interpreted as a qualitative visualization rather than as an independent performance metric.

### 3.4 SHAP-based model interpretation

To interpret how sequence-derived representations and read-mapping-derived quantitative features contributed to ACVD prediction, we performed a unified SHAP analysis for MLMGCN-CVD. Because the DNABERT-S, Nucleotide Transformer, and structural branches generate high-dimensional representations, SHAP values within each branch were aggregated into three modality- level groups. The 17 read-mapping-derived abundance and copy-number features were retained as individual numerical variables. This setting produced an explanation space containing three sequence-derived modality groups and 17 quantitative features, allowing branch-level sequence evidence to be evaluated together with read-mapping-derived abundance and copy-number signals.

Fig. 7A shows the global feature-importance ranking based on mean absolute SHAP values. The three sequence-derived modality groups ranked highest overall. DNABERT-S semantic embedding ranked first, followed by NT contextual embedding and structural representation. Among the read-mapping-derived quantitative features, TPM, mean depth, log_2_(copy number) estimated from mean depth, RPKM, log_2_(copy number) estimated from fragment depth and FPKM ranked higher than the remaining features. These results indicate that MLMGCN-CVD predictions were primarily supported by sequence-derived representations, while read-mapping-derived quantitative variables supplied complementary abundance and copy-number context.

**Fig. 7.**
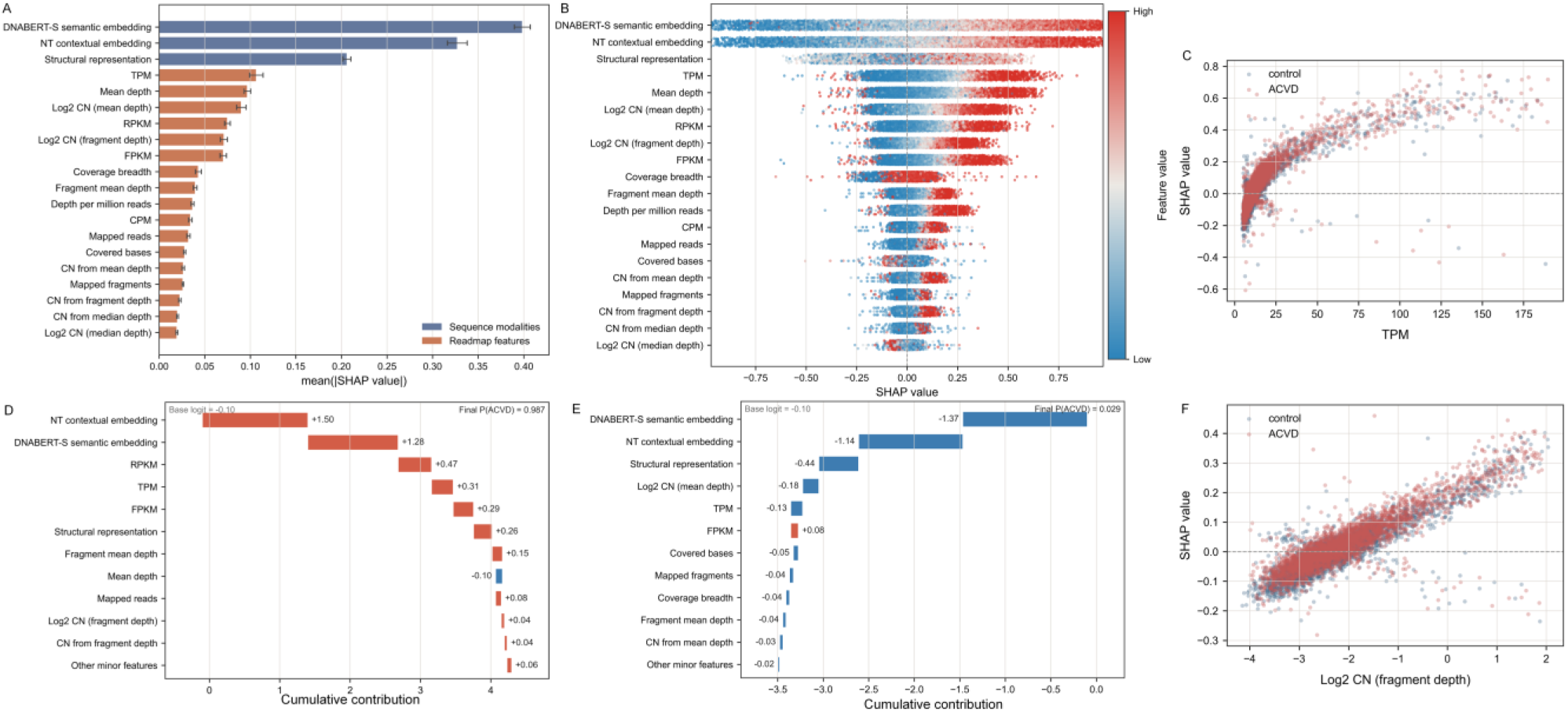
SHAP-based interpretation of sequence-derived modalities and read-mapping-derived quantitative features. (A) Global feature-importance ranking based on mean absolute SHAP values. DNABERT-S, Nucleotide Transformer, and structural representations were aggregated as modality-level groups, whereas the 17 read-mapping-derived quantitative features were retained as individual variables. (B) SHAP summary plot showing the distribution, direction, and feature- value dependence of SHAP contributions. Positive SHAP values indicate increased support for ACVD prediction, whereas negative SHAP values indicate support for the control class. (C) SHAP dependence plot for TPM, showing the relationship between fragment-associated abundance signal and model contribution. (D) Representative waterfall plot for an ACVD-supporting prediction. (E) Representative waterfall plot for a control-supporting prediction. (F) SHAP dependence plot for log_2_(copy number) estimated from fragment depth. This analysis evaluates branch-level and quantitative-feature contributions to model prediction.

Fig. 7B shows the distribution and direction of SHAP contributions. The three sequence-derived modality groups displayed broad positive SHAP ranges. Several read-mapping-derived features, including TPM, mean depth, RPKM and FPKM, also showed positive SHAP contributions in part of the samples. The dependence plot in Fig. 7C showed a nonlinear positive association between TPM and SHAP values, and Fig. 7F showed an increasing trend for log_2_(copy number) estimated from fragment depth.

The representative waterfall plots in Fig. 7D and Fig. 7E showed individual prediction examples. In the ACVD-supporting example, NT contextual embedding, DNABERT-S semantic embedding, RPKM, TPM, FPKM and structural representation contributed positively, resulting in a final ACVD probability of 0.987. In the control-supporting example, DNABERT-S semantic embedding, NT contextual embedding, structural representation, log_2_(copy number) estimated from mean depth and TPM contributed negatively, resulting in a final ACVD probability of 0.029.

### 3.5 Functional annotation and clinical relevance of model high-contribution fragments

To evaluate the biological interpretability of MLMGCN-CVD, we systematically annotated sequence fragments with high model contribution scores. Rather than relying on conventional genus-level abundance comparisons, the analysis focused directly on model-prioritized fragments and examined whether they were enriched in coding regions, recapitulated functional signals previously associated with ACVD, or contained potential regulatory and non-coding RNA elements not previously examined at the fragment level.

High-contribution fragments identified by leave-one-out perturbation analysis were integrated with ORF/CDS prediction using MetaProdigal [48, 49], homologous protein annotation using DIAMOND[50, 51], KEGG functional classification[52], and Infernal/Rfam-based RNA family identification[53, 54]. Enrichment patterns were evaluated using count-based, FPKM-weighted, and copy-number-based strategies. Representative coding fragments and Rfam-supported RNA families were subsequently quantified across samples and correlated with host clinical indices using Spearman correlation analysis.

The majority of high-contribution fragments were located in coding regions, providing a basis for the downstream functional annotation analyses, and the annotation composition and raw recovery counts of high-contribution fragments were also summarized (Fig. S4). High-contribution fragments spanned a broad range of functional categories with heterogeneous enrichment patterns across different quantification strategies (Fig. 8A). Transport and membrane exchange, amino acid/nitrogen metabolism, two-component regulation, DNA replication and repair, and cell envelope/glycan biosynthesis showed case-biased enrichment under at least one analysis setting, whereas RNA translation and methylation, redox and anaerobic respiration, and carbohydrate/energy metabolism tended toward weaker or control-biased enrichment. The connecting lines in Fig. S4 indicate differences in the numbers of case- supporting and control-supporting fragments within each functional category, revealing that most categories contain fragments from both directions with varying degrees of imbalance. This heterogeneity indicates that the model captures functional signals across a broad range of microbial metabolic processes rather than being dominated by a single pathway.

**Fig. 8.**
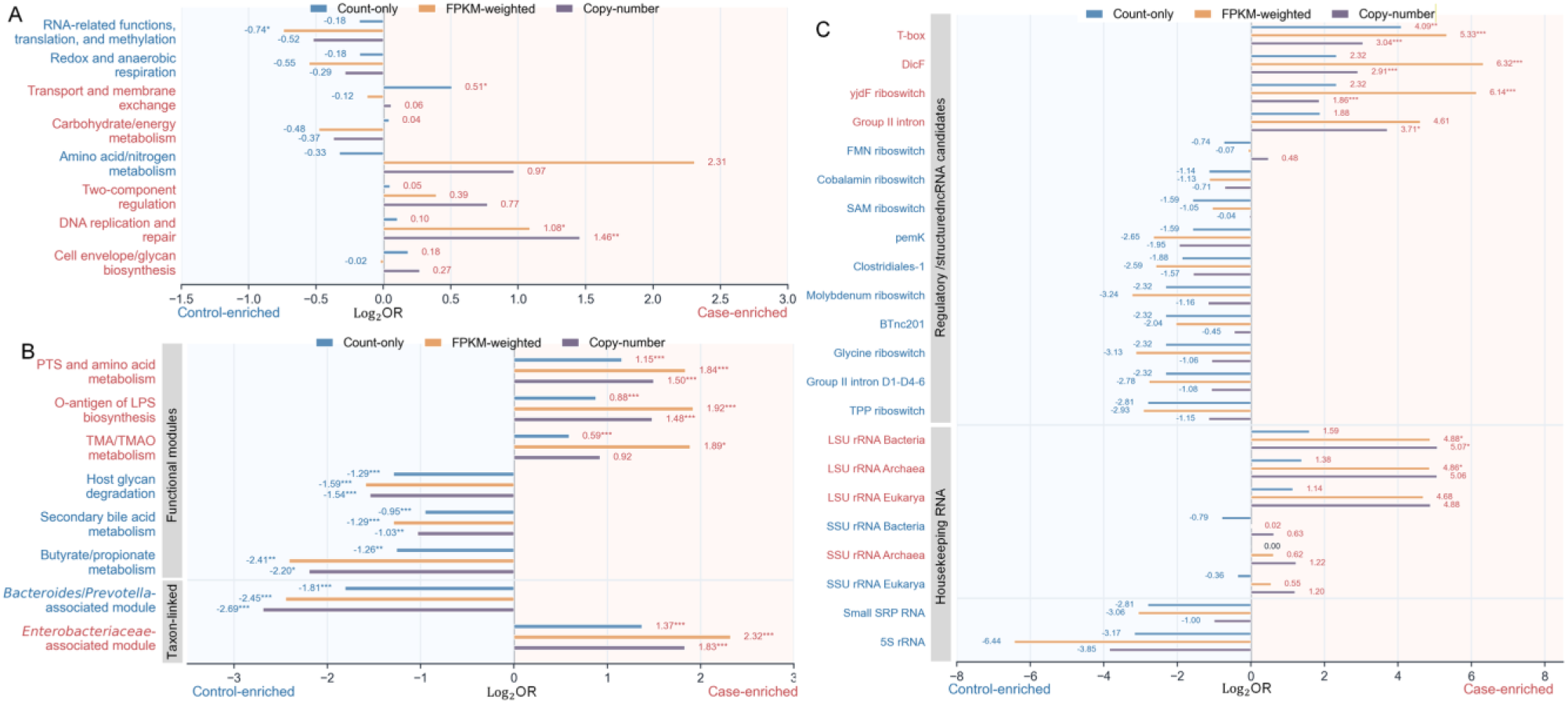
Functional enrichment and RNA-element recovery among model high-contribution fragments. (A) Enrichment patterns of major functional categories among high-contribution fragments. (B) Recovery analysis of ACVD-related functional modules and taxon-linked modules. (C) Enrichment patterns of Rfam-supported regulatory/structured ncRNA candidates and housekeeping RNA families. Bars represent log2 odds ratios calculated under three signal definitions: count-only, FPKM-weighted and copy-number-based analyses. Positive log2OR values indicate case-enriched signals, whereas negative values indicate control-enriched signals. Bar colors denote the three analysis modes: blue, count-only; orange, FPKM-weighted; purple, copy-number. Red and blue row labels indicate the model support direction, with red denoting case-supporting features and blue denoting control-supporting features. Shaded background regions mark the case-enriched and control-enriched sides of the log2OR scale. Grey side labels in panels B and C indicate functional-module or RNA-family groupings. Asterisks denote BH-FDR-adjusted significance levels: * p < 0.05, ** p < 0.01 and *** p < 0.001. OR, odds ratio.

To determine whether the high-contribution fragment-based MLMGCN-CVD could recover functional or taxonomic-background modules previously reported in ACVD metagenomic studies[11], a fragment-level recovery analysis was performed (Fig. 8B). Directional enrichment patterns were broadly consistent across count-only, FPKM-weighted, and copy-number-based analyses, indicating robustness to the quantification strategy. PTS (phosphotransferase system) and amino acid metabolism, LPS/O-antigen biosynthesis, TMA/TMAO metabolism, and the *Enterobacteriaceae*-associated module all showed case-enriched patterns. These findings directly recapitulate the major ACVD-associated functional signals reported by Jie et al.[11] using a metagenome-wide association study (MWAS) approach on the same cohort: enrichment of PTS transport modules, LPS O-antigen biosynthesis (attributed to *Enterobacteriaceae* expansion), TMA lyase genes (YeaW/X), and *Enterobacteriaceae*-associated taxa including *Escherichia coli*, *Klebsiella* spp., and *Enterobacter aerogenes*. In contrast, host glycan degradation, secondary bile acid metabolism, butyrate/propionate metabolism, and the *Bacteroides/Prevotella*-associated module all showed control-enriched patterns. These findings are likewise consistent with Jie et al.[11], who reported reduced potential for glycosaminoglycan degradation, secondary bile acid biosynthesis, SCFA (butyrate and propionate) production in ACVD, as well as *Roseburia intestinalis*, *Faecalibacterium prausnitzii*, *Bacteroides* spp., and *Prevotella copri*. Taken together, the fragment-level recovery analysis demonstrates that MLMGCN- CVD, operating directly on raw sequence fragments without prior taxonomic annotation, independently recapitulates the major functional disease signals identified by the annotation-dependent MWAS approach of Jie et al.[11]. This cross-validation at the functional level provides strong evidence that the model has learned biologically meaningful representations of ACVD-associated microbial activity.

Beyond the recovery of previously reported functional modules, Rfam analysis identified a previously unreported layer of regulatory RNA elements among high-contribution fragments (Fig. 8C). T-box, DicF, yjdF riboswitch, group II intron, and LSU rRNA (Bacteria, Archaea, Eukarya) showed case-enriched patterns, whereas cobalamin riboswitch, SAM riboswitch, pemK, clostridiales-1, molybdenum riboswitch, BTnc201, glycine riboswitch, group II intron D1-D4-6, TPP riboswitch, small SRPRNA, and 5S rRNA showed control-enriched patterns. The TPP, cobalamin, and FMN riboswitches — among the most prevalent and conserved regulatory RNAs in the human gut microbiome — extend the pathway-level observation of reduced vitamin biosynthesis in the ACVD gut microbiome to the regulatory RNA level[55, 56]. Loss of riboswitch-mediated regulatory capacity, rather than merely reduced gene abundance, likely underlies the vitamin biosynthesis deficit. This is compounded by the depletion of *Bacteroides* species capable of TPP riboswitch-regulated thiamine (vitamin B1) biosynthesis and the concurrent enrichment of *Enterobacteriaceae*, whose LPS further suppresses intestinal thiamine absorption and reduces colonic TPP uptake[57, 58]. Cobalamin (vitamin B12) is required for methionine synthase and methylmalonyl-CoA mutase[59]; elevated homocysteine resulting from impaired cobalamin-dependent metabolism is considered a cardiovascular risk factor[60]. Notably, TPP also acts as a potent antagonist of the macrophage P2Y6 receptor, which drives foam cell formation during atherosclerosis; TPP administration reduced atherosclerotic plaque burden in mouse models[61, 62].

The case-enriched DicF sRNA, encoded by the Qin cryptic prophage of *E. coli* [63], aligns with the known *Enterobacteriaceae* enrichment in ACVD[11]. T-box riboswitches, monitoring tRNA aminoacylation status in Gram-positive Bacteria [64, 65], indicate heightened amino acid starvation signaling consistent with concurrent enrichment of amino acid transport, and may partly reflect the expansion of gram-positive *Streptococcus* spp. in the ACVD gut[11]. Collectively, these findings demonstrate that MLMGCN-CVD captures a dimension of ACVD-associated gut microbiome biology — the regulatory RNA landscape — that is not accessible through conventional annotation-dependent approaches.

The case-enriched DicF sRNA, encoded by the Qin cryptic prophage of *E.* fragments, we further evaluated whether representative high-contribution coding fragments were associated with host clinical phenotypes. Candidate fragments were used as reference sequences, and dehosted clean reads from each sample were remapped to these fragments. The abundance-derived signal was represented by log_2_(FPKM + 1), and the copy-number-derived signal was represented by log_2_(copy number). Spearman correlation analysis was then performed for clinical indices related to body composition, blood pressure, lipid metabolism, glucose metabolism, liver function, and kidney function. Representative high-contribution coding fragments displayed heterogeneous clinical correlation patterns across this broad panel of clinical indices (Fig. S5). Case-supporting and control-supporting fragments showed distinct correlation structures, consistent with their opposing roles in ACVD discrimination. Notably, the abundance-derived and copy-number-derived heatmaps were not fully concordant: the ΔAbd/ΔCN panel further showed that some fragment-clinical associations differed between the two quantification backgrounds, indicating that fragment abundance and copy-number background provide non-redundant biological information.

We next examined whether RNA-element families detected among high-contribution fragments were associated with clinical indices. Spearman correlation analysis between Rfam-supported RNA family signals and 13 clinical indices revealed a total of 30 significant associations (p < 0.05) across abundance-derived and copy-number-derived signals, spanning riboswitch/ncRNA, sRNA, housekeeping RNA, and other RNA categories (Fig. 9). Among RNA elements showing consistent correlations across both abundance and copy-number signals, the SAM riboswitch and tmRNA were positively associated with waist-to-hip ratio and systolic BP, respectively. The FMN riboswitch and pemK were each negatively associated with APOA. Positive correlations with APOB were observed for C4 antisense RNA and TwoAYGGAY RNA, whereas DicF sRNA was positively associated with TRIG. Several RNA elements correlated with LDLC: Group II intron, clostridiales-1 RNA, 5S RNA, and TwoAYGGAY RNA showed positive associations, whereas AsdA antisense RNA showed a negative one. C-di-GMP-1 riboswitch and TwoAYGGAY were positively correlated with CHOL. By contrast, group-II-D1D4-5, Bacterial LSU rRNA, and Bacterial large SRP RNA were negatively associated with FBG, and Bacterial RNase P class A showed a negative correlation with CREA.

**Fig. 9.**
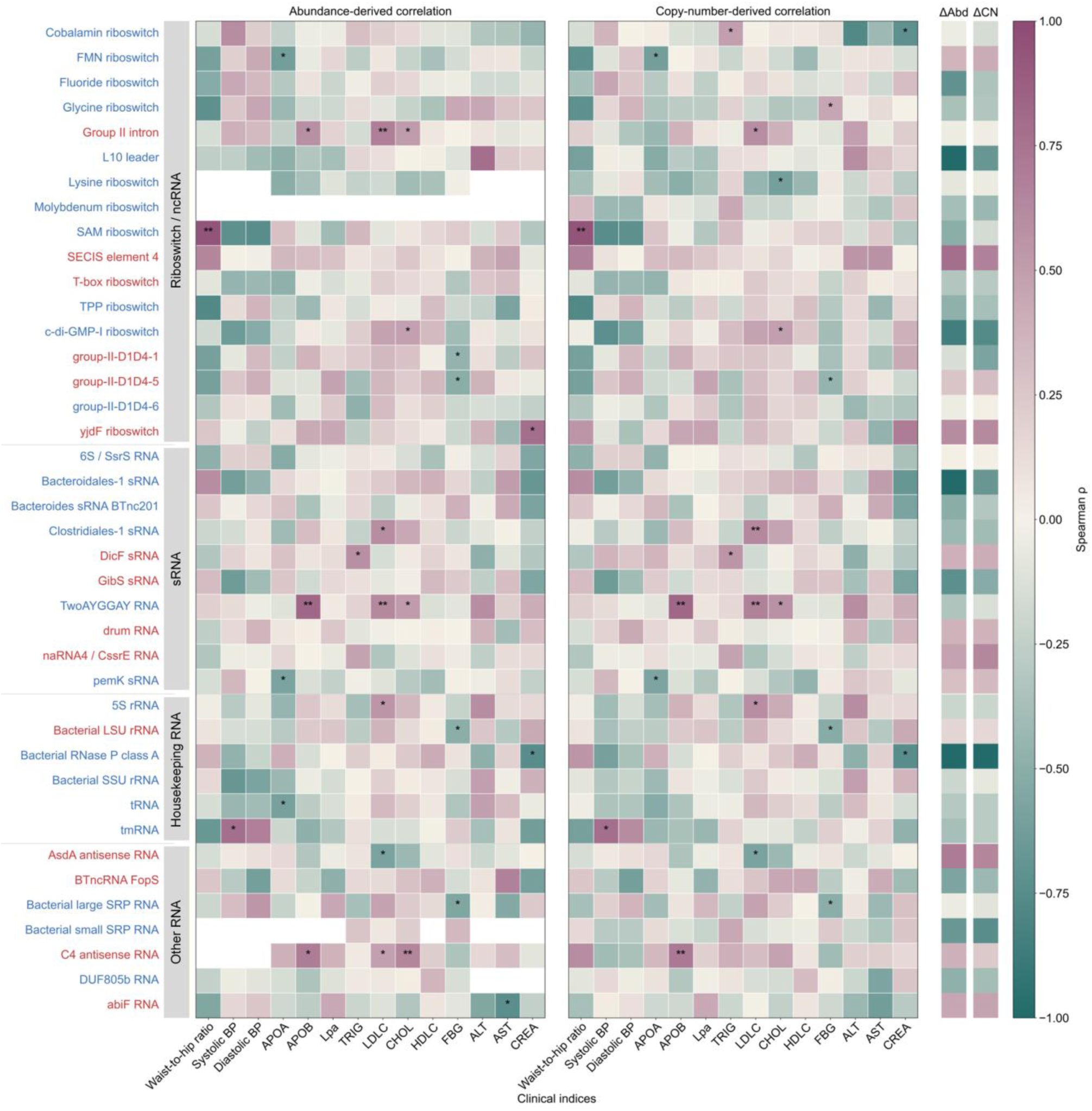
Correlations between Rfam-supported RNA families and clinical indices. Rows represent Rfam RNA families detected in model high-contribution fragments. Grey side labels group RNA families into riboswitch/ncRNA, sRNA, housekeeping RNA and other RNA categories. Red and blue row labels indicate RNA families associated with ACVD-supporting and control-supporting fragment sets, respectively. The left and right heatmaps show Spearman correlations of clinical indices with abundance-derived RNA-family signals and copy-number-derived RNA-family signals, respectively. The heatmap color scale represents Spearman’s ρ, with purple indicating positive correlations, green indicating negative correlations and near-white indicating weak or no correlation. Asterisks denote BH-FDR-adjusted significance. BP, blood pressure; APOA, apolipoprotein A; APOB, apolipoprotein B; Lpa, lipoprotein(a); TRIG, triglycerides; LDLC, low-density lipoprotein cholesterol; CHOL, total cholesterol; HDLC, high-density lipoprotein cholesterol; FBG, fasting blood glucose; ALT, alanine aminotransferase; AST, aspartate aminotransferase; CREA, creatinine. Asterisks denote BH-FDR-adjusted significance levels: * p < 0.05, ** p < 0.01 and *** p < 0.001.

The positive correlation of the SAM riboswitch with waist-to-hip ratio is mechanistically plausible. SAM riboswitches sense intracellular S-adenosylmethionine (SAM) to regulate methionine cycle gene expression[66], and elevated SAM has been associated with fat mass and truncal adiposity in human cohorts[67]. Gut microbiota are major contributors to methionine metabolism[68], and weight gain in obesity has been associated with methionine metabolism perturbations[69], which could be driven by dysbiosis. The positive correlations of Group II introns, 5S rRNA, Clostridiales-1, and TwoAYGGAY with LDLC, and of c-di-GMP-I riboswitch and TwoAYGGAY with CHOL, may shed a light in the mechanism of the established role of gut microbiota in cholesterol metabolism via bile acid biotransformation and short-chain fatty acid production[70]. The negative correlations of housekeeping RNAs (LSU rRNA, RNase P) with FBG and CREA may reflect reduced overall microbial biomass or ribosomal activity in individuals with impaired glucose regulation and renal function, consistent with the known reduction in gut microbial diversity in type 2 diabetes and chronic kidney disease[71].

Jie et al.[11] reported analogous MLG-level correlations with a clinical index panel; the present study extends these findings to the fragment level with higher-resolution associations. These clinical index- associated RNA family signals represent a potentially cost- effective layer of functional biomarkers. The consistent directionality of associations across both abundance-derived and copy-number- derived signals for several RNA families, suggests their potential utility as metagenome-based diagnostic indicators for stratifying cardiometabolic risk in ACVD patients.

## 4 Discussion and Conclusions

This study developed MLMGCN-CVD as a sequence-level multimodal graph-learning framework for ACVD prediction from gut metagenomic sequences without prior taxonomic or functional annotation. Current microbiome disease-prediction frameworks predominantly rely on abundance-based representations derived from annotation-dependent pipelines, including taxonomic profiles, pathway abundance tables, and predefined functional categories. While these approaches have substantially advanced microbiome research, they have several important limitations. First, they inevitably compress heterogeneous metagenomic information into predefined biological units, potentially weaken local sequence variation, regulatory elements, uncharacterized microbial fragments, and latent structural features embedded within metagenomic DNA[11, 13, 14]. Second, they are inherently dependent on the completeness of reference databases: reads from uncharacterized or novel microbial species, which may constitute a substantial fraction of gut microbial diversity, are either discarded or misassigned, potentially losing biologically relevant signals[72, 73]. Third, aggregation to genus or species level discards within-taxon functional heterogeneity: strains of the same species can differ markedly in functional capacity[74]. Yet current shotgun metagenomic sequencing, particularly shallow sequencing (≤1 Gb), can only reliably resolve taxa at the species level[75]. MLMGCN-CVD represents a methodological shift of ACVD prediction from community-level feature engineering toward annotation-free sequence-level representation learning. By directly modelling standardized metagenomic DNA fragments and integrating semantic, contextual, structural and read-mapping-derived information, the framework preserves fine- grained sequence evidence while retaining quantitative support from metagenomic reads. Importantly, the recovery analysis demonstrated that this annotation-free strategy independently recapitulated major ACVD-associated microbial functions previously identified by annotation-dependent metagenome-wide association studies, including TMA/TMAO metabolism, LPS and O-antigen biosynthesis, phosphotransferase systems, and *Enterobacteriaceae*-associated modules. This finding indicates that biologically meaningful disease-associated signals can be learned directly from metagenomic sequences without requiring prior aggregation into predefined taxonomic or pathway categories.

The multimodal embedding analysis further demonstrated that sequence-level microbiome prediction benefits from integrating complementary biological representations. Among the tested modalities, DNABERT-S semantic embeddings contributed the strongest single-modality signal, consistent with the ability of genomic foundation models to capture transferable nucleotide-level regularities through large-scale self-supervised pretraining[35]. Contextual embeddings and sequence-derived structural priors showed weaker independent performance but substantially improved prediction when integrated within the adaptive fusion framework. Importantly, adaptive multimodal fusion outperformed direct feature concatenation, suggesting that performance gains arise not from simple feature accumulation, but from coordinated alignment of heterogeneous biological representations. This observation is particularly relevant for metagenomic disease prediction, where semantic, contextual, structural, and quantitative features differ substantially in dimensionality, scale, and noise characteristics[76].

The backbone ablation experiments further support the importance of multi-scale sequence modelling in metagenomic disease prediction. The superior performance of the Post-CNN configuration suggests that graph propagation first organizes long-range semantic dependencies among related sequence regions, after which convolutional refinement can more effectively capture local motif- like and structure-associated patterns. Biologically, this is consistent with the hierarchical organization of regulatory sequence information, in which local sequence motifs operate within broader contextual and structural environments. More broadly, these findings suggest that sequence-level microbiome signals are inherently relational and multi-scale, supporting the use of graph-learning strategies for modelling interactions among metagenomic fragments[43, 77].

One particularly notable finding was the recovery of regulatory RNA-associated signals among model high-contribution fragments (Fig. 8, Fig. 9, Fig. S5). Although previous ACVD microbiome studies primarily focused on taxonomic abundance shifts and pathway-level alterations, the present work suggests that disease-associated microbiome information may also reside within regulatory sequence architectures. We identified multiple Rfam-supported riboswitches and small regulatory RNAs, including TPP, cobalamin, FMN, glycine, DicF, and pemK-associated elements, several of which showed significant correlations with host cardiometabolic indicators. Importantly, these findings should not be interpreted as direct evidence of causal regulatory activity. Rather, they suggest that regulatory RNA-associated sequence patterns may represent an underexplored layer of microbiome-associated disease information that is largely inaccessible to conventional abundance-based workflows.

Among these signals, vitamin-associated riboswitches displayed particularly interesting enrichment patterns (Fig. 8). TPP, cobalamin, and FMN riboswitch-associated fragments showed control-enriched distributions, extending previous pathway-level observations of reduced vitamin biosynthesis potential in the ACVD microbiome to the regulatory RNA level[11]. Riboswitches are metabolite-responsive regulatory elements that couple intracellular metabolic states to transcriptional or translational control. Their depletion may therefore reflect not only reduced biosynthetic capacity, but also altered regulatory organization within the ACVD- associated gut microbiome. Conversely, the case-enriched DicF small RNA signal was consistent with the expansion of *Enterobacteriaceae*-associated stress-response states previously reported in ACVD cohorts[11]. Together, these observations suggest that microbiome-associated cardiovascular signals may partially reside within regulatory and stress-response architectures that are not readily captured through conventional abundance aggregation.

The external validation results provide preliminary evidence that sequence-level representation learning may improve cross- cohort robustness relative to abundance-based frameworks (Fig. 6). Microbiome prediction models frequently suffer from substantial performance degradation across cohorts because taxonomic abundance profiles are highly sensitive to sequencing depth, preprocessing procedures, population background, and reference-database composition. In contrast, sequence-level representations learned directly from metagenomic fragments may capture more transferable biological regularities that are less dependent on cohort-specific compositional structure. Nevertheless, the present external validation remains limited to a single independent cohort, and broader multicohort evaluation across geographically and technically heterogeneous datasets will be necessary to determine the generalizability of the framework.

The present work also contributes to the emerging field of annotation-free metagenomic deep learning. Conventional microbiome disease-prediction pipelines remain fundamentally constrained by reference-database completeness and taxonomic annotation accuracy. Reads originating from uncharacterized organisms or previously unknown genomic regions are frequently discarded or poorly represented in abundance-based analyses, potentially excluding biologically relevant disease-associated information. By operating directly on standardized metagenomic DNA fragments, MLMGCN-CVD can identify discriminative sequence patterns independently of prior annotation, allowing potentially informative fragments from uncharacterized microbial organisms to contribute to disease modelling. This characteristic may become increasingly important as microbiome studies continue to reveal extensive unexplored microbial genomic diversity.

Several limitations of this study should be acknowledged. First, although the identified RNA-family-associated signals are biologically intriguing, the present work remains computational and does not experimentally validate their regulatory activity or causal contribution to ACVD pathogenesis. Second, the folding-derived structural features used in the ssDNA branch should be interpreted as computational conformational priors rather than direct measurements of in vivo RNA secondary structures. Third, the identified high- contribution fragments represent model-discriminative sequence features and may not necessarily correspond to causal biological determinants. Fourth, despite external validation, the primary cohort was derived predominantly from a single population, and broader validation across ethnic groups, geographic regions, sequencing platforms, and clinical settings will be required before clinical translation. Finally, the observed clinical correlations remain cross-sectional and observational, and future prospective and mechanistic studies will be required to evaluate the longitudinal stability, disease specificity, and functional relevance of the identified sequence elements.

In conclusion, MLMGCN-CVD provides an annotation-free sequence-level framework for microbiome-based ACVD prediction directly from gut metagenomic sequences. By integrating semantic, contextual, structural, and quantitative representations within a unified graph-learning architecture, the framework improves predictive performance while preserving biologically informative local sequence signals. Beyond prediction performance itself, our findings suggest that disease-associated microbiome information extends beyond conventional taxonomic and pathway abundance summaries and may remain partially embedded within raw metagenomic sequences. The recovery of regulatory RNA-associated signals further raises the possibility that regulatory microbiome architectures represent an underexplored dimension of host–microbiome interaction biology. More broadly, this work supports the emerging view that sequence-level representation learning may provide a complementary paradigm for investigating microbiome-associated disease mechanisms and developing next-generation microbiome-based diagnostic frameworks.

## Declarations

### Ethics approval and consent to participate

This study did not involve the recruitment of new human participants, the collection of new human specimens, or the generation of new human-derived sequencing data. All analyses were performed using publicly available, de-identified metagenomic sequencing data and associated metadata from previously published studies. The primary ACVD cohort used in this study was originally approved by the Medical Ethical Review Committee of the Guangdong General Hospital and the Institutional Review Board at BGI-Shenzhen, and informed consent was obtained from all participants, as reported in the original publication by Jie et al.

### Consent for publication

Not applicable.

### Availability of data and materials

The MLMGCN-CVD web server is available at http://www.combio-lezhang.online/MLMGCN-CVD/home. Source code, data, and related reproducibility resources are available at Zenodo under DOI 10.5281/zenodo.18447118 and at GitHub: https://github.com/xing1999/MLMGCN-CVD. The raw sequencing data for the primary cohort are available from the European Nucleotide Archive under accession ERP023788, study PRJEB21528. The external validation cohort data are available from the NCBI Sequence Read Archive under BioProject accession PRJNA615842.

### Authors’ contributions

W.H. conceived the study, developed the methodology, performed the software implementation, curated the data, conducted the formal analysis, prepared the visualizations, and drafted the manuscript. W.W. contributed to study design and methodology, and participated in manuscript writing and revision. L.Z. supervised the study, administered the project, acquired funding, contributed to methodology, and revised the manuscript. Y.V.F. contributed to study design, supervised the study, and revised the manuscript. All authors read and approved the final manuscript.

## Acknowledgements

The authors gratefully acknowledge the financial support from the funding agencies listed in the Funding section.

## Supplementary Information

**Supplementary Methods**

**Method S1. Adaptive semantic graph based on multimodal embeddings Method S2. Deep-learning model framework**

**Supplementary Results**

**Result S1. Comparison with conventional machine-learning baselines**

**Supplementary Figures**

**Figure S1. Adaptive Semantic Graph Modelling Framework Based on Multimodal Embedding. Figure S2. Schematic diagram of the single-strand structural embedding workflow.**

**Figure S3. Performance Comparison Between Deep Learning Models and Traditional Machine Learning Algorithms.**

**Figure S4. Annotation composition and raw recovery counts of model high-contribution fragments.**

**Figure S5. Correlations between representative high-contribution fragments and clinical indices.**

**Supplementary Tables**

**Table S1. Pairwise Wilcoxon Tests on 10-fold Cross-Validation Results Across Feature Modes Table S2. Pairwise Wilcoxon Tests on 10-fold Cross-Validation Results (Post-CNN vs Pre-CNN) Table S3. Pairwise Wilcoxon Tests on 10-fold Cross-Validation Results for Dual-view and Screening-Review**

**Table S4. Pairwise Wilcoxon tests on external-cohort results on PRJNA615842**

