## Supplementary Methods, Tables and Figures for "Annotation-free sequence-level multimodal graph learning reveals gut microbiome signatures of atherosclerotic cardiovascular disease"

### Supplementary Information

#### SUPPLEMENTARY INFORMATION

#### Supplementary Methods

**Method S1. Adaptive semantic graph based on multimodal embeddings**
**Method S2. Deep-learning model framework**

#### Supplementary Results

**Result S1. Comparison with conventional machine-learning baselines**

#### Supplementary Figures

**Figure S1. Adaptive Semantic Graph Modeling Framework Based on Multimodal Embedding.**
**Figure S2. Schematic diagram of the single-strand structural embedding workflow.**
**Figure S3. Performance Comparison Between Deep Learning Models and Traditional Machine Learning Algorithms.**
**Figure S4. Annotation composition and raw recovery counts of model high-contribution fragments.**
**Figure S5. Correlations between representative high-contribution fragments and clinical indices.**

#### Supplementary Tables

**Table S1. Pairwise Wilcoxon Tests on 10-fold Cross-Validation Results Across Feature Modes**
**Table S2. Pairwise Wilcoxon Tests on 10-fold Cross-Validation Results (Post-CNN vs Pre-CNN)**
**Table S3. Pairwise Wilcoxon Tests on 10-fold Cross-Validation Results for Dual-view and Screening-Review**
**Table S4. Pairwise Wilcoxon tests on external-cohort results on PRJNA615842**

### Supplementary Methods

#### Method S1 Adaptive semantic graph based on multimodal embeddings

In recent years, with the broad adoption of deep learning in bioinformatics, it has become increasingly clear that genomic sequences encode not only the linear arrangement of nucleotides but also complex contextual dependencies and structural priors linked to three-dimensional conformations[1, 2]. Conventional sequence encodings(such as one-hot representations, k-mer frequencies, or handcrafted statistical features) achieved early success, yet they typically provide static, local descriptions and therefore struggle to capture long-range dependencies between sequence segments and the underlying regulatory regularities[3]. Inspired by pretrained language models in natural language processing, Transformer-based sequence embedding approaches have been introduced into tasks such as regulatory element identification and modification-site prediction, substantially improving semantic-level representation learning[4]. However, existing methods often prioritise unimodal semantic signals or rely on simple concatenation of multi-source features, with limited systematic consideration of structural/geometric constraints. Moreover, many approaches model sequences purely as linear strings and do not explicitly characterise non-local interactions among segments, which constrains their capacity to represent higher-order biological features and cross-segment organisational patterns[5].

To address these limitations, we propose an Adaptive Semantic Graph Modelling (ASGM) framework built on multimodal embeddings, which reconstructs the sequence representation space from three complementary perspectives: semantics, long-range context, and structure. ASGM first integrates pretrained language models with computable structural priors to extract complementary features at the semantic level (DNABERT-S)[6], long-range contextual level (Nucleotide Transformer), and structural level (DNAshape or RNAfold/RNAplfold). These modality-specific features are then aligned into a shared latent space via linear projection and normalisation, followed by adaptive fusion using attention- and/or gating-based mechanisms. Finally, ASGM computes inter-segment semantic similarity in the fused space to construct a content-aware graph, enabling the model to jointly optimise node representations and graph connectivity during learning and to obtain a context-dependent dynamic graph representation. This design not only strengthens the expressive power of multimodal fusion, but also provides an explicit structural formulation of latent interactions, yielding a semantically coherent and biologically plausible topological prior for higher-order information propagation in downstream graph learning.

##### Method S1.1 DNABERT-S semantic embedding

Within the multimodal embedding framework, the semantic modality aims to learn contextual dependencies and semantic regularities in DNA sequences, thereby revealing latent functional patterns. In this study, we employ DNABERT-S (a genome-scale pretrained language model) as the semantic feature extractor to obtain contextual embeddings with cross-species generalisation. Built on a bidirectional Transformer architecture, DNABERT-S treats DNA as a biological language and is pretrained on multi-species genomes using masked language modelling and curriculum-based contrastive learning, yielding a high-dimensional representation space that captures long-range dependencies and semantic consistency. Given a DNA sequence $S = \{b_1, b_2, \dots, b_N\}$, the model first tokenises the sequence into fixed-length $k$-mers:

$$x_i = (b_i, b_{i+1}, \dots, b_{i+k-1}),\ i = 1, \dots, L \quad (1)$$

Each token is mapped to a dense vector $E(x_i) \in \mathbb{R}^d$ via an embedding layer. The token embeddings are then combined with positional encodings $P_i$, and then they were fed into a multi-layer bidirectional Transformer to capture global contextual dependencies among nucleotide segments:

$$\begin{cases} H^{(l)} = \text{Transformer}^{(l)}(H^{(l-1)}) \\ H^{(0)} = [E(x_1) + P_1, \dots, E(x_L) + P_L] \end{cases} \quad (2)$$

where $H^{(l)}$ denotes the output representation of the $l$-th Transformer layer. Context-aware dependency modelling is achieved via the self-attention mechanism using query ($Q$), key ($K$), and value ($V$) projections:

$$\text{Attention}(Q, K, V) = \text{softmax}\left(\frac{QK^\top}{\sqrt{d_k}}\right)V \quad (3)$$

which enables each $k$-mer representation to aggregate information from other segments in the sequence according to semantic relevance. DNABERT-S is optimised with a masked language modelling objective, maximising the reconstruction probability of masked tokens conditioned on the remaining context:

$$\mathcal{L}_{\mathrm{MLM}} = -\sum_{i\in\mathcal{M}} \log P(x_i \mid x_{\setminus\mathcal{M}}) \tag{4}$$

where $M$ denotes the set of randomly masked tokens. To further regularise the global structure of the embedding space, DNABERT-S incorporates contrastive learning at higher-level semantic representations, with the objective:

$$\mathcal{L}_{\mathrm{CL}} = -\frac{1}{2B}\sum_{i=1}^{B} \log \frac{\exp\left(s(f(x_i), f(x_i^{+}))/\tau\right)}{\sum_{j\neq i} \alpha_{ij} \exp\left(s(f(x_i), f(x_j))/\tau\right)} \tag{5}$$

where $s(\cdot,\cdot)$ is the cosine similarity, $\tau$ is a temperature parameter, and $\alpha_{ij}$is a similarity-based weighting term. After semantic encoding, we obtain the token-level representation $H = [h_1, h_2, \dots, h_L]$. A sequence-level semantic vector is then derived by mean pooling:

$$z_{\mathrm{DNABERT-S}} = \frac{1}{L}\sum_{i=1}^{L} h_i \tag{6}$$

The resulting vector $z_{\text{DNABERT-S}} \in \mathbb{R}^{d_1}$ integrates local nucleotide semantics with global contextual dependencies, and is used as the semantic input of the multimodal system together with the contextual and structural modalities for subsequent modality alignment and graph construction.

**Method S1.2 Nucleotide Transformer contextual embedding**

To further increase the model's capacity to capture global dependencies, we introduce the Nucleotide Transformer (NT) as a contextual feature extractor in addition to the semantic embeddings. By modelling long-range context, NT learns semantic coherence and distal dependencies within DNA sequences. Given an input sequence $S = \{b_1, b_2, \dots, b_N\}$, the sequence is similarly tokenised into a set of fixed-window $k$-mers $X = \{x_1, x_2, \dots, x_L\}$. During embedding, each token is projected into a continuous space:

$$\mathrm{e}_i = \mathrm{W}_E \phi(x_i) + \mathrm{p}_i \tag{7}$$

where $\phi(\cdot)$denotes an index-based encoding, $W_E$is the embedding matrix, and $p_i$is the positional vector. Through multi-layer contextual modelling, NT can adaptively disentangle contributions from local versus long-range dependencies, yielding a more robust global contextual representation. The final layer outputs a contextual representation sequence:

$$H^{(L)} = \{h_1^{(L)}, h_2^{(L)}, \dots, h_L^{(L)}\} \tag{8}$$

which is then aggregated into a global contextual vector via attention-weighted pooling:

$$z_{\mathrm{NT}} = \sum_{i=1}^{L} \omega_i h_i^{(L)}, \omega_i = \frac{\exp(\beta_i)}{\sum_j \exp(\beta_j)} \tag{9}$$

where $\omega_i$is an attention-derived weight that quantifies each segment's contribution to the global semantics. This contextual embedding provides a global semantic prior for subsequent modality alignment and graph construction, complementing the semantic modality by strengthening long-range dependency modelling and enabling multi-scale information fusion.

**Method S1.3 Double-stranded structural embedding**

Beyond semantic and contextual sequence representations, the local geometry of DNA plays a critical role in protein binding, chromatin accessibility, and gene regulation. The three-dimensional conformation of DNA is not solely determined by nucleotide order; it is reflected in physical properties including variations in minor-groove width, helical twisting, and local bending. These factors provide structural constraints that complement linear "sequence semantics" in mapping sequence to function. To characterise double-stranded conformational features, we introduce DNAshapeR as the structural modality to quantify local geometric attributes of the DNA double helix, thereby complementing the semantic and contextual modalities. Based on statistical mechanics and molecular-dynamics-derived models, DNAshapeR parameterises local DNA geometry using four core descriptors: Helix Twist (HelT), Minor Groove Width (MGW), Propeller Twist (ProT), and Roll. For a given sequence $S = \{b_1, \dots, b_N\}$, nucleotide-resolution structural features are computed as

$$\mathrm{S}_i = [\mathrm{HelT}_i, \mathrm{MGW}_i, \mathrm{ProT}_i, \mathrm{Roll}_i] \tag{10}$$

And it yields a structural matrix $S \in \mathbb{R}^{L\times 4}$. To increase the expressive capacity of structural information, we further apply a lightweight encoder to perform contextual modelling over the DNAshape feature sequence. Specifically, each position-wise structural vector is first projected onto a $d_s$-dimensional space:

$$\mathrm{h}_i^{(0)} = \mathrm{W}_s \mathrm{s}_i + \mathrm{b}_s \tag{11}$$

and subsequently updated by a structure-aware encoding module:

$$\mathrm{h}_i^{(l)} = \mathrm{LN}\left(\mathrm{h}_i^{(l-1)} + \mathrm{FFN}(\mathrm{Attn}(\mathrm{h}_i^{(l-1)}))\right) \tag{12}$$

Finally, a sequence-level structural embedding is obtained via mean pooling:

$$z_{\mathrm{Shape}} = \frac{1}{L}\sum_{i=1}^{L} \mathrm{h}_i^{(L)} \tag{13}$$

Through this DNAshape channel, our model incorporates computable geometric priors that promote biological plausibility when learning regulatory patterns, while providing complementary information to the semantic and contextual modalities. An overview of the dsDNA-model ASGM pipeline (spanning DNABERT-S semantic embedding, NT contextual embedding, DNAshape double-stranded geometric embedding, multimodal fusion, and semantic graph construction) is illustrated by Fig. S1.

**Method S1.4 Single-stranded structural embedding**

In the ssDNA-model, the structural modality no longer relies on the double-stranded geometric descriptors provided by DNAshape. Instead, the input sequence is treated as a single-stranded molecule, and secondary structure is inferred by RNAfold and RNAplfold, from which statistical features are extracted to capture structural cues such as single-strand folding patterns and local accessibility. This design is intended to approximate structural states that may arise under local strand separation, transient single-stranded exposure, or the involvement of ssDNA-associated components, thereby enabling the structural channel to provide information complementary to double-stranded geometry.

Specifically, RNAfold is used to compute the secondary structure corresponding to the minimum free energy (MFE) conformation. From the predicted structure, we derive features that reflect the global folding state and stability, including the sequence-level free energy, the fractions of paired and unpaired bases, length statistics of structural elements (e.g., stems and loops), and the base-pairing status sequence obtained from the dot–bracket representation. In addition, RNAplfold provides nucleotide-resolution statistics such as local base-pairing probabilities and unpaired probabilities, which characterise the propensity of local regions to form stable pairings and their accessibility, thereby mitigating uncertainty that may arise when relying solely on a single MFE structure.

To maintain consistency with the semantic and contextual modalities at the node granularity, we aggregate nucleotide-level structural features to $k$-mer nodes. Concretely, we apply a sliding-window mean pooling over the structural feature sequence by $k$-mers ($k = 6$), yielding node-level structural vectors. In the ssDNA-model, these node-level structural vectors replace the DNAshapeR geometric descriptors used in the dsDNA-model, while all other components (modality alignment and fusion, semantic graph construction, and the downstream graph-learning backbone) remain unchanged. Through this "structural-channel substitution" strategy, the ssDNA-model incorporates a single-stranded secondary-structure prior without altering the overall framework, forming a structural-view complement to the dsDNA-model. Here, the complete pipeline is illustrated by Fig. S2.

**Method S1.5 Multimodal alignment and construction of a semantic-fusion graph**

After obtaining the semantic feature $z_{\text{DNABERT-S}}$, the contextual feature $z_{\text{NT}}$, and the structural feature (either $z_{\text{Shape}}$for dsDNA or $z_{\text{SS}}$for ssDNA), we design a multimodal alignment and semantic fusion mechanism, as well as construct a semantics-driven and content-aware graph in the fused space. The goal is to integrate heterogeneous information across modalities so that the model can represent multi-level properties of DNA sequences in a unified latent space, thereby providing structured inputs for higher-order feature propagation in downstream graph neural networks. First, embeddings from these three modalities are mapped onto a shared latent space via modality-specific projection layers:

$$\tilde{z}_m = \mathrm{W}_m z_m + \mathrm{b}_m, m \in \{\text{BERT-S, NT, Shape}\} \tag{14}$$

where $W_m$is a learnable linear transformation that aligns scale and distributional differences across modalities. Next, a crossmodal attention/gating mechanism is used to learn modality weights and perform fusion:

$$z_{\text{fusion}} = \sum_m \alpha_m \tilde{z}_m \tag{15}$$

where $\alpha_m$ quantifies the contribution of each modality for the current sequence, enabling the fused representation to adaptively balance semantic, contextual, and structural information. Based on the fused embeddings, we then construct a content-aware semantic graph to capture semantic dependencies among sequence segments. Nodes correspond to $k$-mer derived segments, and edges are formed according to inter-segment semantic similarity. Specifically, we connect each node to its $K$ nearest neighbours in the similarity space. Let $h_i$ and $h_j$ denote the fused features of the $i^{\text{th}}$ and $j^{\text{th}}$ segments, respectively. Their similarity is defined as cosine similarity:

$$A_{ij} = \sin(h_i, h_j) = \frac{h_i^{\top} h_j}{\|h_i\| \|h_j\|} \tag{16}$$

To prevent overly dense graphs, we keep the Top-$K$ neighbours for each node and introduce a self-loop term $\epsilon$ to improve the stability of message passing. The adjacency matrix is thus defined as:

$$A_{ij} = \begin{cases} \sin(h_i, h_j), & j \in \mathcal{N}_K(i) \\ \epsilon, & i = j \\ 0, & \text{otherwise} \end{cases} \tag{17}$$

In practice, the neighborhood relations can be initialized by an attention-weight matrix from DNABERT-S as a prior, and then be updated by the fused representations, allowing the graph structure to reflect both local semantic dependence and cross-segment contextual association. In this way, each DNA sequence is represented as a semantic-aware graph:

$$\mathcal{G} = (\mathcal{V}, \mathcal{E}, A) \tag{18}$$

This graph preserves local contextual relations while explicitly modelling potential long-range dependencies at the global level, providing a differentiable and interpretable topological prior for subsequent graph learning and bridging multimodal features with structural representations.

#### Method S2. Deep-learning model framework

Metagenomic sequences used for ACVD risk identification often exhibit complex regulatory signals, hierarchical heterogeneity, and cross-scale dependencies. Building upon the multimodal feature embeddings and content-aware semantic graph constructed in the preceding section, we further develop a hierarchical deep-learning framework that combines a graph convolutional network with a one-dimensional convolutional neural network, termed MLMGCN-CVD, to jointly model topological dependencies among sequence segments and local motif or structural patterns. The model takes the semantic graph G=(V,E,A) generated by ASGM as the main input, where node features are the aligned and fused representations from DNABERT-S, Nucleotide Transformer, and the structural channel. The structural channel is instantiated as dsDNA geometric features from DNAshapeR in the dsDNA-Model and as ssDNA folding statistics from RNAfold or RNAplfold in the ssDNA-Model. Edge weights are adaptively determined by semantic similarity in the fused space to capture latent inter-segment associations. In addition to this graph-based sequence representation, MLMGCN-CVD incorporates readmap-derived abundance and copy-number features as auxiliary global quantitative features. These features are not used for node construction or graph construction, because they describe fragment-level read-mapping support and copy-number background rather than position-specific token information. Instead, they are introduced through a projection-and-gating branch at the final late-fusion stage. Overall, MLMGCN-CVD consists of a deep graph convolutional backbone, a convolutional feature refinement layer, a multi-head attention readout, a quantitative feature projection and gating branch, and a late-fusion classifier. This hierarchical design enables progressive information aggregation from local to global in the semantic-graph space and combines sequence-level representations with quantitative read-mapping context for ACVD discrimination.

**(1) Deep graph convolutional backbone**

The model receives the semantic graph $G = (V, E, A)$ as input. To stabilise feature propagation, we add self-loops and apply symmetric normalisation to the adjacency matrix:

$$\tilde{A} = A + I, \quad \hat{A} = \tilde{D}^{-\frac{1}{2}} \tilde{A} \tilde{D}^{-\frac{1}{2}} \tag{19}$$

where $\tilde{D}$ is the degree matrix of $\tilde{A}$. Let $H^{(l)}$ denote node representations at layer $l$. The graph convolution update is:

$$H^{(l)} = \sigma\left(\hat{A}H^{(l-1)}W^{(l)}\right) + H^{(l-1)} \tag{20}$$

where $W^{(l)}$ is a learnable weight matrix, $\sigma(\cdot)$ is the ReLU activation, and the residual connection alleviates over-smoothing in deep graph networks and increases training stability. In addition, each graph convolution block can incorporate a lightweight channel-reweighting module to adaptively emphasize discriminative semantic and structural cues across channels.

**(2) Convolutional feature refinement layer**

Neighbourhood aggregation alone may be insufficient to capture stable local patterns, such as short motifs and fine-grained conformational cues. Therefore, after obtaining the graph-convolution output $H^{(L)}$, we apply a one-dimensional convolutional refinement module along the node dimension to enhance local patterns within a limited receptive field:

$$H_{\text{cnn}} = \text{ConvRefine1D}(H^{(L)}) \tag{21}$$

In implementation, this module uses convolutional transformation, nonlinear activation, dropout, normalization, and residual updating to refine node-level features without changing the hidden dimension. This module preserves the global propagation capacity of graph learning, while explicitly extracting salient local segment patterns and complementing the topological information encoded by the graph.

**(3) Multi-head attention readout**

Next, node-level representations are aggregated into a sequence-level vector. We adopt a multi-head attention readout to perform weighted pooling over nodes. For the $t$th attention head, the aggregated representation is:

$$r_t = \sum_{i\in\mathcal{V}} \beta_i^{(t)} h_i, \beta_i^{(t)} = \frac{\exp(\mathrm{q}_t^{\mathrm{T}} h_i)}{\sum_j \exp(\mathrm{q}_t^{\mathrm{T}} h_j)} \tag{22}$$

where $h_i$ denotes the input node representation from $H_{\text{cnn}}$, and $q_t$ is the learnable query vector for head $t$. Outputs from all heads are concatenated and linearly mapped to obtain the graph-derived sequence representation:

$$z_{\text{graph}} = \text{ReLU}(\mathrm{W}_r[r_1 \parallel r_2 \parallel \cdots \parallel r_T]) \tag{23}$$

This readout mechanism selects discriminative segments across multiple attention subspaces, enabling effective compression and integration of global sequence information.

**(4) Late-fusion classifier and loss function**

In addition to the graph-derived representation $z_{\text{graph}}$, MLMGCN-CVD uses two global auxiliary branches before final classification. The first branch processes the sequence-level Nucleotide Transformer representation $u_{\text{NT}}$. It projects $u_{\text{NT}}$ into the hidden space to obtain $z_{\text{seq}}$, and generates a gate vector $\gamma_{\text{seq}}$ to modulate the graph-derived representation:

$$z_{\text{seq}} = f_{\text{seq}}(u_{\text{NT}}),\ \gamma_{\text{seq}} = f_{\text{gate}}(u_{\text{NT}}) \tag{24}$$

The second branch processes the readmap-derived abundance and copy-number feature vector $x_{\text{quant}}$. This quantitative branch maps the numerical vector into the hidden representation space and generates a quantitative gate:

$$z_{\text{quant}} = f_{\text{quant}}(x_{\text{quant}}), \gamma_{\text{quant}} = f_{\text{qgate}}(x_{\text{quant}}) \tag{25}$$

Here, $z_{\text{quant}}$ provides an explicit representation of fragment-level abundance and copy-number context, whereas $\gamma_{\text{quant}}$ modulates the graph-derived sequence representation according to read-mapping support. The final fused representation is defined as:

$$z_{\text{fused}} = [z_{\text{graph}} \odot (1 + \gamma_{\text{seq}} + \gamma_{\text{quant}}) \parallel z_{\text{seq}} \parallel z_{\text{quant}}] \tag{26}$$

where $\odot$ denotes element-wise multiplication and $\parallel$ denotes vector concatenation. In our implementation, $z_{\text{graph}}$, $z_{\text{seq}}$, and $z_{\text{quant}}$ are all mapped to the same hidden dimension. With a hidden dimension of 768, the fused representation is 2304-dimensional before entering the final classifier. The predicted probability of ACVD is then obtained by:

$$\hat{y} = \sigma(\text{MLP}(z_{\text{fused}})) \tag{27}$$

During training, we use a class-weighted binary cross-entropy loss to mitigate class imbalance:

$$L = -w_1 y \log \hat{y} - w_0(1-y)\log(1-\hat{y}) \tag{28}$$

where $w_1$ and $w_0$ are the positive and negative class weights, respectively, and $y \in \{0,1\}$ is the ground-truth label.

In summary, MLMGCN-CVD takes the content-aware semantic graph constructed by ASGM as the main input, models inter-segment topological dependencies via deep graph convolution, refines local motif and structural patterns via a one-dimensional convolutional refinement module, and obtains a graph-derived sequence representation by multi-head attention readout. Readmap-derived abundance and copy-number features are incorporated as auxiliary global quantitative features through a late-fusion projection and gating branch rather than being injected into node-level inputs. This design jointly captures sequence-level semantic evidence, structural topology, local patterns, and fragment-level quantitative context, enabling more effective modelling of cross-scale signals in metagenomic sequences and increasing prediction stability. For the ssDNA-Model, the network architecture and training objective remain unchanged; only the structural channel of the input node features is replaced, using single-stranded secondary-structure statistics inferred by RNAfold or RNAplfold instead of double-stranded geometric features from DNAshape. The readmap-derived quantitative branch is kept unchanged across the dsDNA-Model and ssDNA-Model.

### Supplementary Results

#### Result S1. Comparison with conventional machine-learning baselines

To further establish a comparable reference between our method and the conventional machine-learning pipelines commonly used in prior studies, we conducted a supplementary benchmark against classic tabular-learning baselines. In many microbiome-based disease risk prediction studies, a typical paradigm is to first represent each sample as a feature vector and then feed it into a conventional classifier for discrimination. Following this protocol, we used the multimodal fused feature representation from the main text as a unified input, namely the MLMFusion representation. The machine-learning baselines included five representative algorithms: Random Forest (RF), Logistic Regression (LR), k-Nearest Neighbors (KNN), Support Vector Machine (SVM), and Extreme Gradient Boosting (XGBoost). All models were trained and evaluated under the same stratified 10-fold cross-validation framework as in the main text. This supplementary comparison therefore provides a direct performance reference under identical inputs and an identical evaluation protocol, enabling us to assess the additional gains and result stability brought by deep structured modeling. The results are reported in Fig. S3.

Fig. S3 presents the comparative results between MLMGCN-CVD and five conventional machine-learning baselines under the same fused input representation. As shown in Fig. S3A, MLMGCN-CVD achieves the highest values in both sensitivity and specificity, indicating a more balanced prediction performance for ACVD and control samples. The fold-wise distributions in Fig. S3B and Fig. S3C further show that MLMGCN-CVD obtains the highest ACC and AUC among all compared models. Compared with RF, KNN, SVM, LR, and XGBoost, the performance distribution of MLMGCN-CVD is shifted toward a higher value range, and the significance annotations indicate clear improvements over the conventional baselines. Fig. S3D and Fig. S3E show consistent trends in F1-score and MCC, where MLMGCN-CVD achieves the highest mean values and maintains a relatively concentrated distribution across folds. These results suggest that the proposed model not only improves prediction accuracy, but also provides more stable fold-level performance.

Among the conventional machine-learning methods, XGBoost shows the strongest overall performance, followed by SVM, RF, LR, and KNN in most metrics. However, XGBoost still performs lower than MLMGCN-CVD across ACC, AUC, F1-score, and MCC. The ROC curves in Fig. S3F further confirm this result. MLMGCN-CVD achieves the highest AUC of 0.978, exceeding XGBoost with 0.962, SVM with 0.954, RF with 0.953, LR with 0.947, and KNN with 0.923. Since all compared classifiers use the same MLMFusion representation as input, these results indicate that the improvement is not solely attributable to the fused features. The deep structured modeling framework of MLMGCN-CVD further contributes to performance gains by capturing nonlinear and graph-structured relationships that are difficult for conventional tabular classifiers to fully exploit. Overall, this supplementary benchmark provides additional evidence that MLMGCN-CVD has stronger discriminative ability and higher stability than classical machine-learning pipelines under an identical input and evaluation setting.

### Supplementary Figures

#### Figure S1

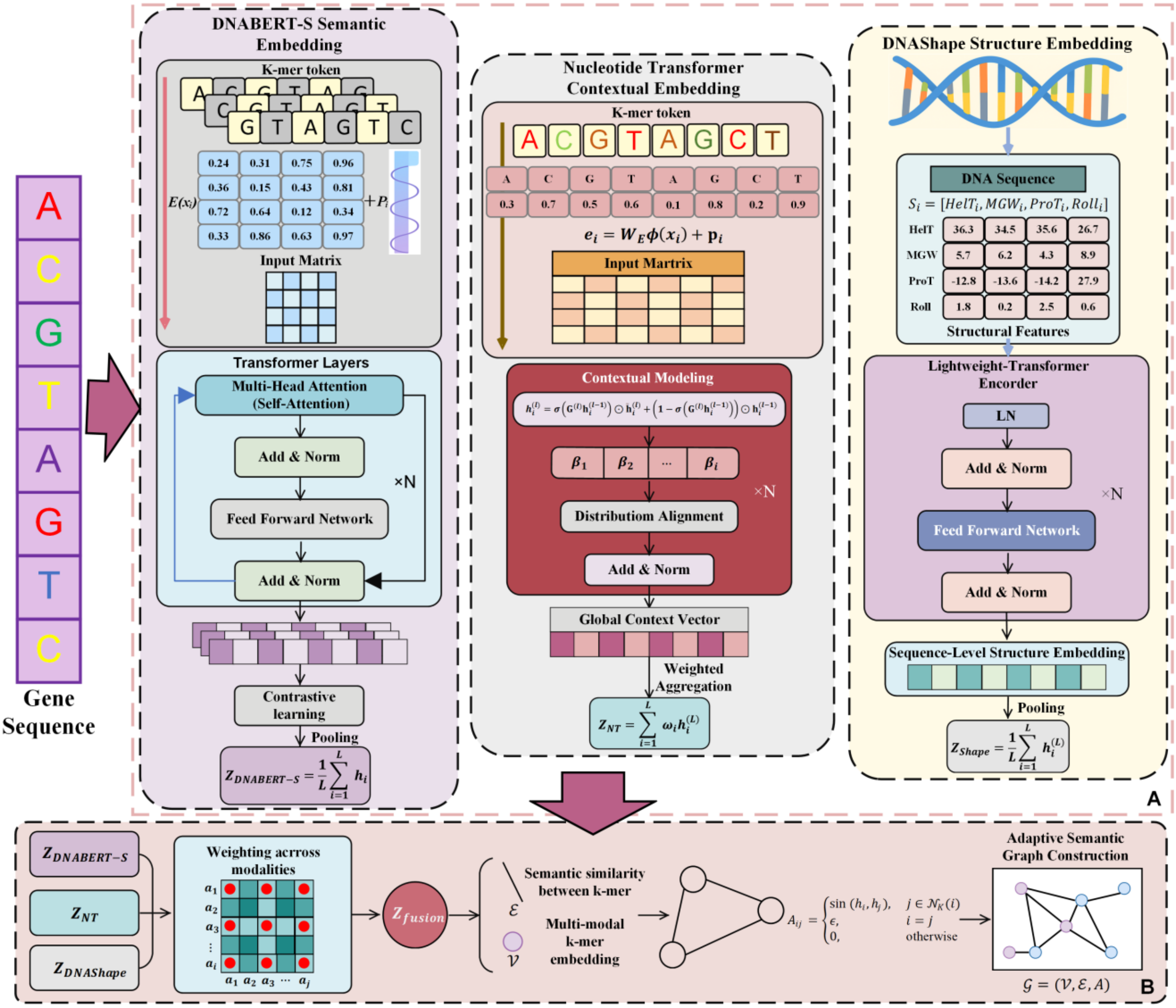


Fig S1. Adaptive Semantic Graph Modeling Framework Based on Multimodal Embedding.

#### Figure S2

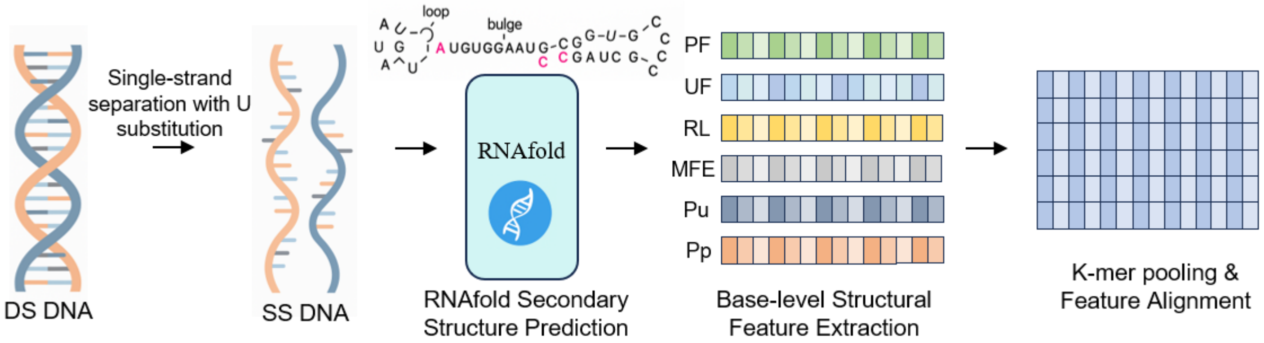


Fig S2. Schematic diagram of the single-strand structural embedding workflow.

#### Figure S3

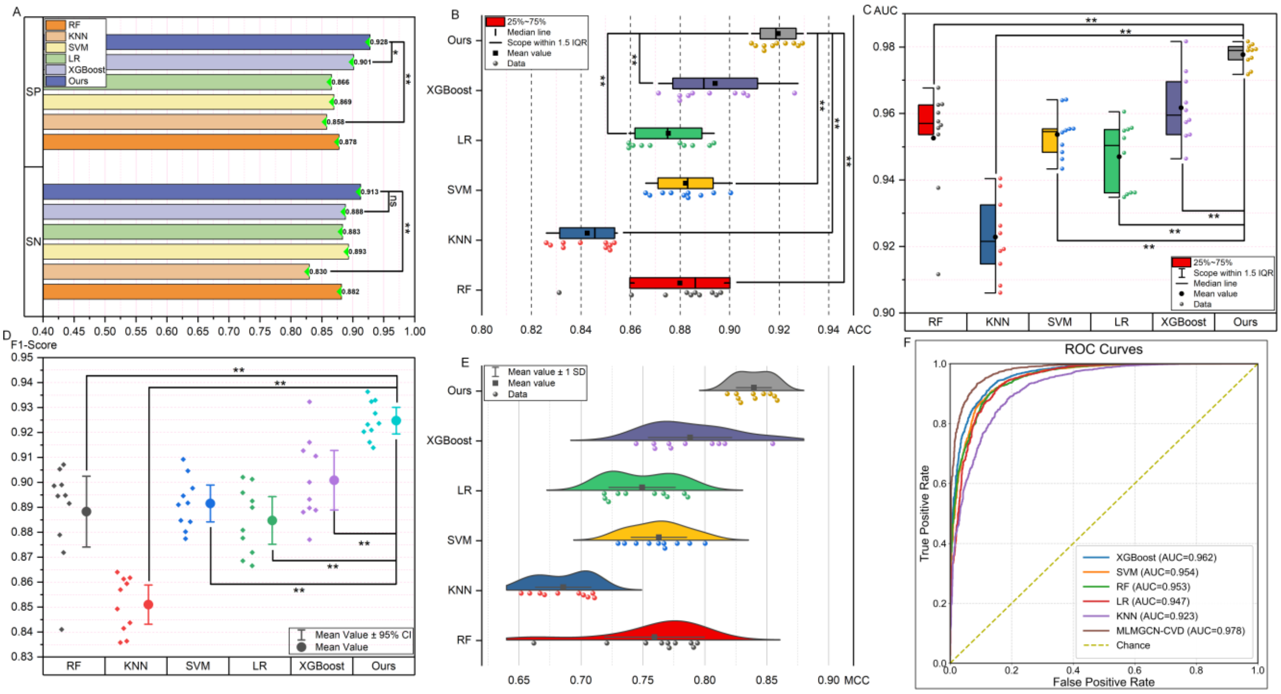


**Fig S3. Performance Comparison Between Deep Learning Models and Traditional Machine Learning Algorithms.** (A) SN and SP comparison across methods. (B) ACC distributions across folds. (C) AUC distributions across folds. (D) F1-score with mean and 95% confidence interval. (E) MCC distributions (fold-wise density). (F) ROC curves of ML baselines vs. ours.

### Figure S4

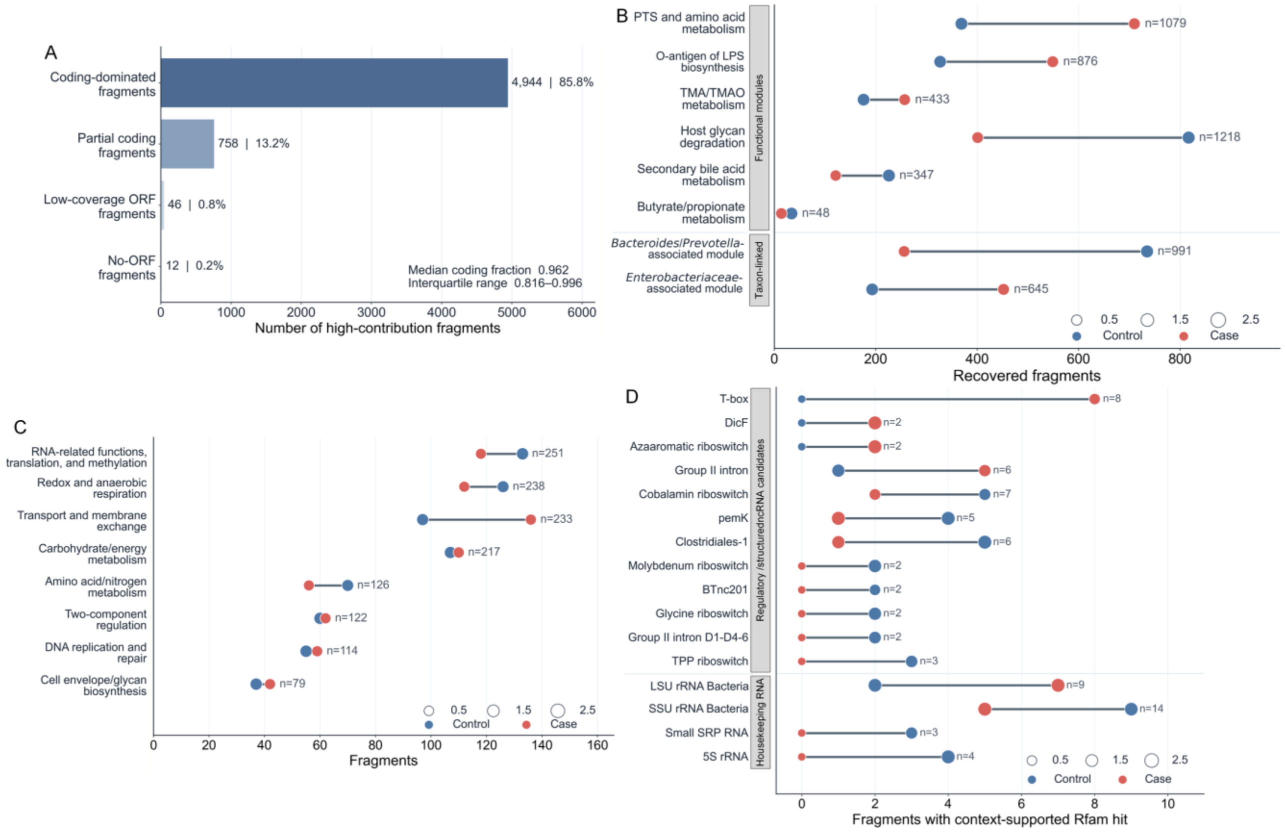


**Fig. S4. Annotation composition and raw recovery counts of model high-contribution fragments.** (A) ORF/CDS-based sequence-region composition of model high-contribution fragments. Numbers and percentages indicate the count and proportion of fragments in each annotation class. The median coding fraction and interquartile range are shown for coding-dominated fragments. (B) Raw recovery counts of previously reported ACVD-associated functional modules and taxon-linked modules among high-contribution fragments. (C) Raw recovery counts of major functional categories detected among high-contribution coding fragments. (D) Raw recovery counts of Rfam-supported RNA-family hits among high-contribution fragments, grouped into regulatory or structured ncRNA candidates and housekeeping RNA families. In panels B–D, blue and red points indicate control-supporting and ACVD-supporting fragments, respectively, and horizontal segments connect the two support directions within each category. The n labels indicate the total number of recovered fragments in each category across both support directions. Point size denotes the absolute enrichment magnitude.

#### Figure S5

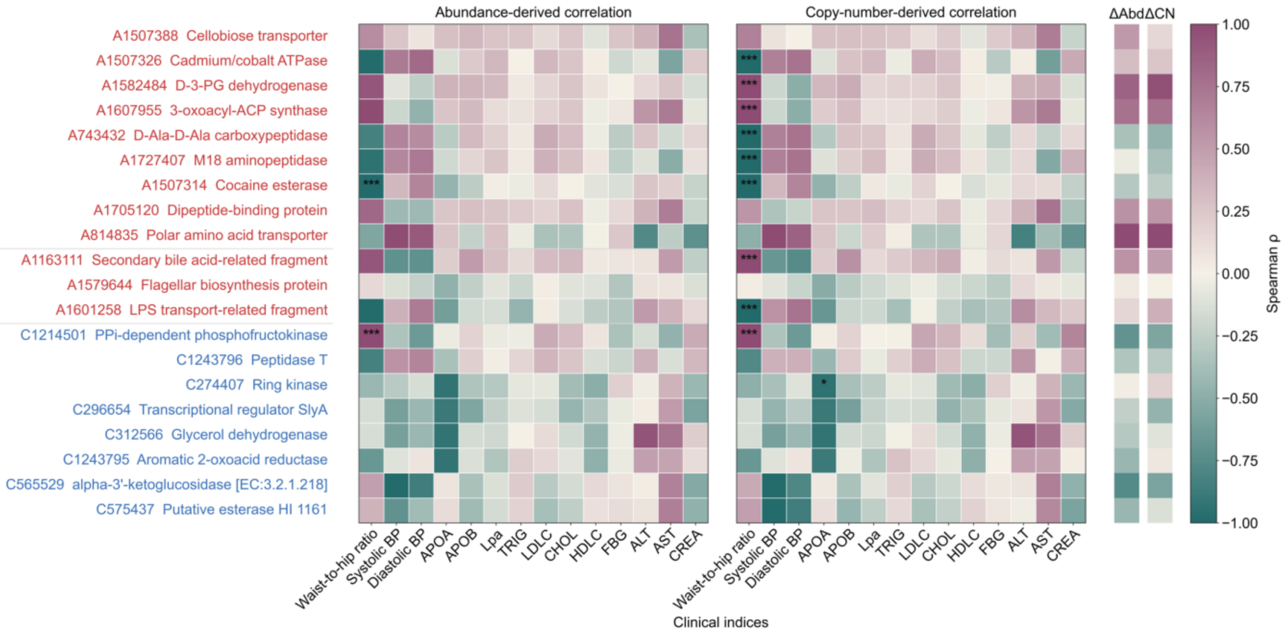


**Fig. S5. Correlations between representative high-contribution fragments and clinical indices.** Rows represent model-prioritized candidate fragments annotated by fragment ID and putative function. Red and blue row labels indicate ACVD-supporting and control-supporting fragments, respectively. The left and middle heatmaps show Spearman correlations between clinical indices and abundance-derived signals, $\log_2(\text{FPKM} + 1)$, or copy-number-derived signals, $\log_2(\text{copy number})$. The heatmap color scale represents Spearman's ρ, with purple indicating positive correlations, green indicating negative correlations and near-white indicating weak or no correlation. Asterisks denote BH-FDR-adjusted significance, * $p < 0.05$, ** $p < 0.01$ and *** $p < 0.001$. The two right-side bars summarize group-difference directions for abundance and copy-number signals, denoted as ΔAbd and ΔCN. BP, blood pressure; APOA, apolipoprotein A; APOB, apolipoprotein B; Lpa, lipoprotein(a); TRIG, triglycerides; LDLC, low-density lipoprotein cholesterol; CHOL, total cholesterol; HDLC, high-density lipoprotein cholesterol; FBG, fasting blood glucose; ALT, alanine aminotransferase; AST, aspartate aminotransferase; CREA, creatinine.

### Supplementary Tables

#### Table S1

**Table S1. Pairwise Wilcoxon Tests on 10-fold Cross-Validation Results Across Feature Modes**

| Comparison | ACC | | | MCC | | |
|---|---|---|---|---|---|---|
| | Δ mean | P-Value | P-FDR(BH) | Δ mean | P-Value | P-FDR(BH) |
| DNABERT-S-only vs NT-only | 0.0366 | 0.001953 | 0.002734 | 0.0784 | 0.001953 | 0.002734 |
| DNABERT-S-only vs Structural-only | 0.0521 | 0.001953 | 0.002734 | 0.1073 | 0.001953 | 0.002734 |
| NT-only vs Structural-only | 0.0155 | 0.083984 | 0.083984 | 0.0289 | 0.130859 | 0.130859 |
| MLMFusion vs Fusion(S+Struct) | 0.1032 | 0.001953 | 0.002734 | 0.202 | 0.001953 | 0.002734 |

| | | | | | | |
|---|---|---|---|---|---|---|
| MLMFusion vs Fusion(Struct+NT) | 0.0911 | 0.001953 | 0.002734 | 0.1795 | 0.001953 | 0.002734 |
| MLMFusion vs Fusion(S+NT) | 0.0549 | 0.001953 | 0.002734 | 0.1111 | 0.001953 | 0.002734 |
| MLMFusion vs Fusion(All,Concat) | 0.0214 | 0.003906 | 0.004557 | 0.0433 | 0.003906 | 0.004557 |

Note: Δmean denotes the average paired difference across the 10 folds (Δmean = mean[score(Model A) - score(Model B)]), where a positive value indicates Model A outperforms Model B. Statistical significance was assessed using a paired Wilcoxon signed-rank test (two-sided) on the per-fold results from sample-level stratified grouped 10-fold cross-validation. P-FDR(BH) indicates p-values adjusted by the Benjamini–Hochberg procedure to control the false discovery rate within each metric across the seven predefined comparisons.

#### Table S2

**Table S2. Pairwise Wilcoxon Tests on 10-fold Cross-Validation Results (Post-CNN vs Pre-CNN)**

| Comparison | ACC | | | MCC | | |
|---|---|---|---|---|---|---|
| | Δ mean | P-Value | P-FDR(BH) | Δ mean | P-Value | P-FDR(BH) |
| Post vs Pre (avg over CNN; GNN_0_Layer) | 0.0239 | 0.001953 | 0.002441 | 0.049 | 0.001953 | 0.002441 |
| Post vs Pre (avg over CNN; GNN_2_Layer) | 0.018 | 0.005859 | 0.005859 | 0.0342 | 0.005859 | 0.005859 |
| Post vs Pre (avg over CNN; GNN_4_Layer) | 0.0201 | 0.003906 | 0.00434 | 0.0392 | 0.003906 | 0.00434 |
| Post vs Pre (avg over CNN; GNN_6_Layer) | 0.0284 | 0.001953 | 0.002441 | 0.0558 | 0.001953 | 0.002441 |
| Post vs Pre (avg over CNN; GNN_8_Layer) | 0.0203 | 0.001953 | 0.002441 | 0.0399 | 0.001953 | 0.002441 |
| Post vs Pre (avg over CNN; GNN_10_Layer) | 0.024 | 0.001953 | 0.002441 | 0.0479 | 0.001953 | 0.002441 |
| Post vs Pre (avg over GNN; CNN_0_Layer) | 0.0229 | 0.001953 | 0.002441 | 0.0449 | 0.001953 | 0.002441 |
| Post vs Pre (avg over GNN; CNN_1_Layer) | 0.0207 | 0.001953 | 0.002441 | 0.0404 | 0.001953 | 0.002441 |
| Post vs Pre (avg over GNN; CNN_2_Layer) | 0.0239 | 0.001953 | 0.002441 | 0.0482 | 0.001953 | 0.002441 |
| Post vs Pre (avg over GNN; CNN_3_Layer) | 0.0223 | 0.001953 | 0.002441 | 0.0439 | 0.001953 | 0.002441 |

Note: Δmean denotes the average paired difference across 10 folds (Δmean = mean[score(Post) - score(Pre)]). P-values are from a two-sided paired Wilcoxon signed-rank test on per-fold results. P-FDR(BH) indicates p-values adjusted by the Benjamini–Hochberg procedure within each metric across the 10 predefined comparisons.

#### Table S3

**Table S3. Pairwise Wilcoxon Tests on 10-fold Cross-Validation Results for Dual-view and Screening-Review**

| Comparison | ACC | | | AUC | | |
|---|---|---|---|---|---|---|
| | Δ mean | P-Value | P-FDR(BH) | Δ mean | P-Value | P-FDR(BH) |
| ssDNA-Model vs dsDNA-Model | 0.0054 | 0.556641 | 0.556641 | 0.0028 | 0.232422 | 0.232422 |
| Screening-Review vs dsDNA-Model | 0.0212 | 0.003906 | 0.005859 | 0.013 | 0.001953 | 0.005859 |
| Screening-Review vs ssDNA-Model | 0.0158 | 0.003906 | 0.005859 | 0.0102 | 0.003906 | 0.005859 |

Note: Δmean denotes the average paired difference across the 10 folds (Δmean = mean[score(Model A) - score(Model B)]). Two-sided paired Wilcoxon signed-rank tests were performed on per-fold results. P-FDR(BH) indicates Benjamini–Hochberg adjusted p-values within each metric across the predefined comparisons.

#### Table S4

**Table S4. Pairwise Wilcoxon tests on external-cohort results on PRJNA615842**

| Comparison | ACC | | | AUC | | |
|---|---|---|---|---|---|---|
| | Δ mean | P-Value | P-FDR(BH) | Δ mean | P-Value | P-FDR(BH) |
| Baseline-Relative-Abundance vs Baseline-Pathway | -0.027296 | 0.013672 | 0.027344 | -0.022491 | 0.027344 | 0.054688 |

| | | | | | | |
|---|---|---|---|---|---|---|
| Baseline-Relative-Abundance vs Sequence-Only | 0.001983 | 0.695313 | 0.695313 | -0.000708 | 1 | 1 |
| Baseline-Relative-Abundance vs MLMGCN-CVD | -0.041465 | 0.003906 | 0.017578 | -0.037276 | 0.001953 | 0.011719 |
| Baseline-Pathway vs Sequence-Only | 0.029279 | 0.048828 | 0.073242 | 0.021784 | 0.105469 | 0.126563 |
| Baseline-Pathway vs MLMGCN-CVD | -0.01417 | 0.232422 | 0.278906 | -0.014784 | 0.105469 | 0.126563 |
| Sequence-Only vs MLMGCN-CVD | -0.043449 | 0.005859 | 0.017578 | -0.036568 | 0.003906 | 0.011719 |

Note: Δmean is the mean paired performance difference, calculated as Δmean = mean[score(Model A) - score(Model B)], where a positive value indicates that Model A outperforms Model B. Two-sided paired Wilcoxon signed-rank tests were performed on the 10 paired results from external evaluation on PRJNA615842. P-FDR(BH) indicates Benjamini–Hochberg adjusted p-values within each metric across the predefined comparisons.
